# Comprehensive evaluation of machine learning models and gene expression signatures for prostate cancer prognosis using large population cohorts

**DOI:** 10.1101/2021.07.02.450975

**Authors:** Ruidong Li, Jianguo Zhu, Wei-De Zhong, Zhenyu Jia

**Affiliations:** Department of Botany and Plant Sciences, University of California, Riverside, CA, USA; Graduate Program in Genetics, Genomics, and Bioinformatics, University of California, Riverside, CA, USA; Department of Urology, Guizhou Provincial People’s Hospital, Guizhou, China; Department of Urology, Guangdong Key Laboratory of Clinical Molecular Medicine and Diagnostics, Guangzhou First People’s Hospital, School of Medicine, South China University of Technology, Guangzhou, China; Urology Key Laboratory of Guangdong Province, The First Affiliated Hospital of Guangzhou Medical University, Guangzhou Medical University, Guangzhou, China; Macau Institute for Applied Research in Medicine and Health, Macau University of Science and Technology, Macau, China

**Author notes:** To whom correspondence should be addressed: Ruidong Li at and Zhenyu Jia at.

**Keywords:** prostate cancer, prognostic signature, machine learning, gene expression, clinical decision-making

## Abstract

Overtreatment remains the pervasive problem in prostate cancer (PCa) management due to the highly variable and often indolent course. Molecular signatures derived from gene expression profiling have played critical roles in PCa treatment decision-making. Many gene expression signatures have been developed to improve the risk stratification of PCa and some of them have already been translationally applied to clinical practice, however, no comprehensive evaluation was performed to compare the performances of the signatures. In this study, we conducted a systematic and unbiased evaluation of 15 machine learning (ML) algorithms and 30 published PCa gene expression-based prognostic signatures leveraging 10 transcriptomics datasets with 1,558 primary PCa patients from public data repositories. The results revealed that survival analysis models outperformed binary classification models for risk assessment, and the performances of the survival analysis methods - Cox model regularized with ridge penalty (Cox-Ridge) and partial least squares regression for Cox model (Cox-PLS) – were generally more robust than the other methods. Based on the Cox-Ridge algorithm, a few top prognostic signatures that have comparable or even better performances than the commercial panels have been identified. The findings from the study will greatly facilitate the identification of existing prognostic signatures that are promising for further validations in prospective studies and promote the development of robust prognostic models to guide clinical decision-making. Moreover, the study provided a valuable data resource from large primary PCa cohorts, which can be used to develop, validate, and evaluate novel statistical methodologies and molecular signatures to improve PCa management.

## Introduction

Prostate cancer (PCa) is the second most frequently diagnosed cancer in men worldwide, with an estimated 1.4 million new cases and 375,000 deaths in 2020 (1). Localized PCa is a highly heterogeneous disease which may lead to variable clinical outcomes. The majority of prostate tumors grow slowly and patients with low-risk PCa may only need active surveillance, whereas patients with aggressive PCa will require immediate treatment. Radical prostatectomy (RP) is the primary treatment for localized PCa with good oncologic outcomes (2). However, approximately 20-40% of patients will experience biochemical recurrence (BCR), i.e., escalated prostate-specific antigen (PSA) levels within 10 years after RP (3–5). Despite of considerable efforts, it remains challenging to accurately predict the aggressiveness of disease at the time of diagnosis to identify patients who only need active surveillance, and to select patients who may benefit from adjuvant treatments, such as chemotherapy, radiation, or immunotherapy following definitive therapy. Thus, overtreatment, which may cause various side effects and impact the quality of patients’ lives, continues to be a pervasive problem in PCa management.

In the past decades, many gene expression-based signatures have been developed for risk assessment in PCa using various computational approaches or statistical methodologies. The aim was to identify a set of cancer-associated genes for constructing a statistical model to predict clinical outcomes. A variety of ML algorithms, which can efficiently handle the high-dimensional transcriptomics data where the number of variables (i.e., genes) is substantially greater than the number of observations (i.e., patients), have been widely used for the development of prognostic models in cancer research. The ML algorithms can be classified into two categories: (i) supervised learning such as least absolute shrinkage and selection operator (LASSO), support vector machines (SVM), and random forest (RF), etc. and (ii) unsupervised learning such as principal component analysis (PCA) and variational autoencoders (VAE), etc. Various strategies have been adapted to model the clinical outcome data. For instance, some of the prognostic models were developed by dividing patients into two risk groups, i.e., aggressive and indolent, to identify signature genes (e.g., differentially expressed genes) to develop a binary classifier (6,7), whereas many other models were built by selecting prognostic genes based on their associations with the time to BCR and modeling the censored time-to-event data for the prediction of relapse-free survival (RFS) (8–10). In addition, unsupervised ML techniques, such as hierarchical clustering, non-negative matrix factorization (NMF) clustering, and latent process decomposition (LPD) were also commonly used to identify cancer subtypes with clinical relevance, and a binary or multi-class classifier will eventually be developed for patient stratification (11–13).

Molecular signatures derived from gene expression profiling have become important risk assessment tools to assist on therapeutic decision-making for PCa, including the identification of candidates for active surveillance, selection of the type and intensity of treatment, and determination of the benefit from adjuvant therapy or multimodal therapy, etc. Some of the published signatures have already been applied to clinical practice, including Decipher (6), Oncotype DX Genomic Prostate Score (GPS) (14), and Prolaris (8). Evaluation of prognostic signatures has been performed in several cancer types such as breast cancer and lung cancer (15–17), however, no systematic and unbiased comparison has been performed to evaluate the performances of multiple ML algorithms and prognostic signatures for PCa. To fill this void, we leveraged ten public transcriptomics datasets consisting of 1,558 primary PCa cases to comprehensively evaluate the performances of 15 ML algorithms and 30 published gene expression signatures for PCa prognosis. Generally, the survival models using the time-to-event (i.e., time to BCR) data outperformed the binary classification models which categorizes patients into “aggressive” (BCR occurred) group and “indolent” (BCR-free) group given 5-year follow-up time. Two survival analysis methods, i.e., Cox-Ridge and Cox-PLS, were more robust than the other methods or algorithms. Most of the 30 prognostic signatures showed certain prognostic powers, while a few signatures, including Penney (18), Wu (10), Li (19), and Sinnott (20), had comparable as or even superior performance than the commercial panels. These promising prognostic signatures, once validated with prospective trials, may be included in clinical practice to improve PCa management. In addition, our study demonstrated that the performances of random gene expression sets (RGESs), the whole transcriptome, and individual genes in the published signatures all ranked lower than the top published signatures, indicating that it’s critical to identify a set of signature genes that are significantly associated with clinical outcomes from the transcriptome to achieve a high level of accuracy for cancer prognosis.

This is the first study that comprehensively evaluated the performances of multiple ML algorithms and published prognostic signatures using PCa population cohorts of large sizes. The findings from the study have demonstrated the necessity and clinical impact of deriving molecular signatures from the transcriptome profiling for PCa prognosis, greatly facilitated the selection of promising prognostic signatures for further validations in prospective clinical studies, and helped to refine the strategy for novel and robust prognostic model development to assist on PCa treatment decision-making. Moreover, we have established a valuable data resource, which consists of 10 transcriptomics datasets with a total of 1,558 PCa cases and a collection of 30 gene expression prognostic signatures, for the development, validation, and evaluation of novel statistical methodologies and molecular signatures to improve PCa management.

## Materials and Methods

### Collection of PCa transcriptomics data from public data repositories

A comprehensive search for transcriptomics data of patients with primary PCa was performed in the public data repositories, including the National Cancer Institute (NCI) Genomic Data Commons (GDC) (21), cBioportal (22), National Center for Biotechnology Information (NCBI) Gene Expression Omnibus (GEO) (23), and ArrayExpress (19). The following criteria were used for dataset selection: (i) the dataset must have a complete record of the time to BCR or the time to the last follow-up if no BCR is incurred after treatment; (ii) the sample size should be greater than 80; and (iii) the dataset must be generated using a genome-wide gene expression profiling platform. A total of 10 datasets were selected in the study and the detailed information were provided in Supplementary Table S1.

The HTSeq-Counts data from The Cancer Genome Atlas Prostate Adenocarcinoma (TCGA-PRAD) project were downloaded and preprocessed using the R package *GDCRNATools* (25). The raw. CEL files of the four Affymetrix microarray datasets including CPC-Gene (GSE107299), Taylor (GSE21034), CancerMap (GSE94764), and CIT (E-MTAB-6128) were downloaded from GEO/ArrayExpress and normalized with the Robust Multichip Average (RMA) method implemented in the R package *oligo* (26). The raw sequencing data for the GSE54460 dataset was downloaded from SRA (https://www.ncbi.nlm.nih.gov/sra) under the accession number SRP036848 using *fasterq-dump* in the SRA Toolkit (version 2.10.8). *STAR* (version 2.7.2a) (27) was used for sequence alignment and *featureCounts* (version 2.0.0) (28) was used for gene expression quantification. The count data was then normalized using the Trimmed Mean of M values (TMM) method implemented in the R package *edgeR* (29). The reads per kilobase per million mapped reads (RPKM) values for the DFKZ dataset was downloaded from cBioPortal and log2 transformation was performed. The processed intensity data of the other datasets including Cambridge (GSE70768), Stockholm (GSE70769), and Belfast (GSE116918) were downloaded directly from GEO using the R package *GEOquery* (30). Gene/probe IDs in all the datasets were converted to the corresponding Ensembl IDs. If multiple probes/genes matched to the same Ensembl ID, only the most informative one with the maximum interquartile range (IQR) for the gene expression is used for this Ensembl ID. The *ExpressionSet* objects of the processed data, including the normalized gene expression data and harmonized metadata, have been deposited in the PCaDB database (31). The gene expression values in each dataset were rescaled by z-score transformation for model evaluation.

### Collection of published gene expression signatures for PCa prognosis

Gene expression-based signatures for PCa prognosis were collected by an inclusive literature screening. The keywords ‘prostate cancer’, ‘prognosis, and ‘gene expression signature’ were used to search the PubMed database. Some signatures were identified in recent review papers on prostate cancer prognostic signatures or in research papers with signature comparisons (32–34). Different types of gene identifiers may be reported in the original papers, so the gene IDs were harmonized by searching against the Ensembl (35), the HUGO Gene Nomenclature Committee (HGNC) (36), and the NCBI Entrez Gene (37) databases. A total of 30 gene expression signatures for PCa prognosis were collected and evaluated in the study (Supplementary Table S2). The gene list of the 30 published prognostic signatures were provided in Supplementary Table S3.

### Binary classification algorithms

Nine binary classification algorithms were evaluated in the study, including Elastic Net, support vector machines (SVM) using the linear function (SVM-Linear), the polynomial kernel function (SVM-Poly), and the radial basis function (SVM-RBF), random forest (RF), partial least square (PLS), linear discriminant analysis (LDA), and XGBoost using the linear booster (XGBoost-Linear) and the tree booster (XGBoost-Tree).

The R package *caret* (38), which has a set of functions to streamline the process for creating predictive models, was used for model training and parameter tuning. In each model, the predictor variables were the genes in a given signature and the response variable was the 5-year BCR status, where 0 represents non-BCR and 1 represents BCR incurred within 5 years after treatment. The grid search with a tuning length of 10 was used for tuning parameters. The 10-fold cross validation resampling scheme and ROC metric were used to select the optimal model. A probability score for the BCR class was calculated for each patient, where a higher score indicates a higher probability to experience BCR.

### Survival analysis methods

Six survival analysis methods were evaluated in the study, including Cox proportional hazards (CoxPH), Cox model regularized with ridge penalty (Cox-Ridge) and lasso penalty (Cox-Lasso), supervised principal components (SuperPC), partial least squares regression for Cox models (Cox-PLS), and random survival forest (RSF).

The CoxPH models were built using the R package *survival (*https://CRAN.R-project.org/package=survival). A linear combination of the expression values and coefficients of the signature genes were computed as the risk scores for the patients. Cox-Ridge and Cox-Lasso models were built using the R package *glmnet* (39) with the penalty α=0 and α=1, respectively. The R package *superpc* (40) was used to build the SuperPC models. A subset of genes in a signature with the univariate regression coefficients exceeding the threshold of 0.3 was used to calculate the principal components to build the model. Cox-PLS models were built using the R package *plsRcox* (41) and two components were included in the models. The R package *randomForestSRC* (42) was used to build the RSF models with 100 trees.

### Comparison of machine learning models and prognostic signatures

#### Intra-dataset (within dataset) comparison

The 10-fold cross-validation (CV) was performed for any dataset to evaluate the performance of a signature using a predictive model within the dataset. In a 10-fold CV, the patient cases were randomly partitioned into ten portions with approximately equal size. In each iteration, nine portions were used as the training set to develop the prediction model, and the risk scores (or class probabilities for binary classification models) were calculated for patients in the remaining one portion (test set). This process was repeated ten times such that each portion was used exactly once as the test data. Identical data partitioning was used to generate 10 folds for evaluating all the prediction models.

#### Inter-dataset (across datasets) comparison

For the inter-dataset comparisons, the prediction models were trained with one dataset and then tested by the other nine independent datasets. Note we have selected ten datasets that met the three selection criteria for this study.

#### Evaluation Metrics

We used three metrics, i.e., concordance index (C-index), time-dependent receiver operating characteristics (ROC) curve, and hazard ratio (HR) estimated by the Kaplan Meier (KM) analysis, to evaluate the performances of the machine learning models and prognostic signatures. The C-index was calculated using the R package *survcomp* (43), the area under the ROC curves (AUC) was estimated using the R package *survivalROC* (44), and KM analyses were performed using the R package *survival* to estimate the HR and 95% confidence intervals (CIs). For the KM analyses, the median values of the risk scores were used to dichotomize the patients into low- and high-risk groups.

### Differential expression analysis, CoxPH survival analysis, and functional enrichment analysis for the signature genes

The R packages *limma* (45) was used to perform differential expression (DE) analysis and *survival* was used for CoxPH survival analysis to investigate whether the genes in the prognostic signatures were differentially expressed between tumor and normal samples, and if they were significantly associated with RFS outcomes based on the TCGA-PRAD data. Kyoto Encyclopedia of Genes and Genomes (KEGG) and Disease ontology (DO) enrichment analysis were performed using the R package *clusterProfiler* (46).

## Results

### Overview of the study design

A total of 50 public transcriptomics datasets of patients with primary PCa were identified from the public data repositories, including GDC, cBioportal, GEO, and ArrayExpress, while only 14 of them have the BCR status data for the patients. Eventually, 10 transcriptomics datasets (a total of 1,558 cases) which met three data selection criteria (*see Methods*) were included in the study. The clinical characteristics of the 10 cohorts were summarized in Supplementary Table S4. We gleaned 30 published gene expression prognostic signatures for PCa through a comprehensive literature search. Nine binary classification algorithms and six survival analysis methods were used to build the prediction models based on the expression data of genes in each prognostic signature. The comprehensive evaluation included comparisons of: (i) the predictive abilities of binary classification models versus survival analysis models as well as between different survival algorithms; (ii) the prognostic performances of the 30 gene expression signatures based on the robust survival algorithm, and (iii) the performances of RGESs, individual signature genes, union of the signature genes, and the whole transcriptome with the published signatures. Both intra-dataset comparison and inter-dataset comparisons were conducted, and three metrics including C-index, time-dependent AUC, and HR based on KM survival analysis were used to evaluate the models and signatures. The overall design of the study was presented in Figure 1.

**Figure 1.**
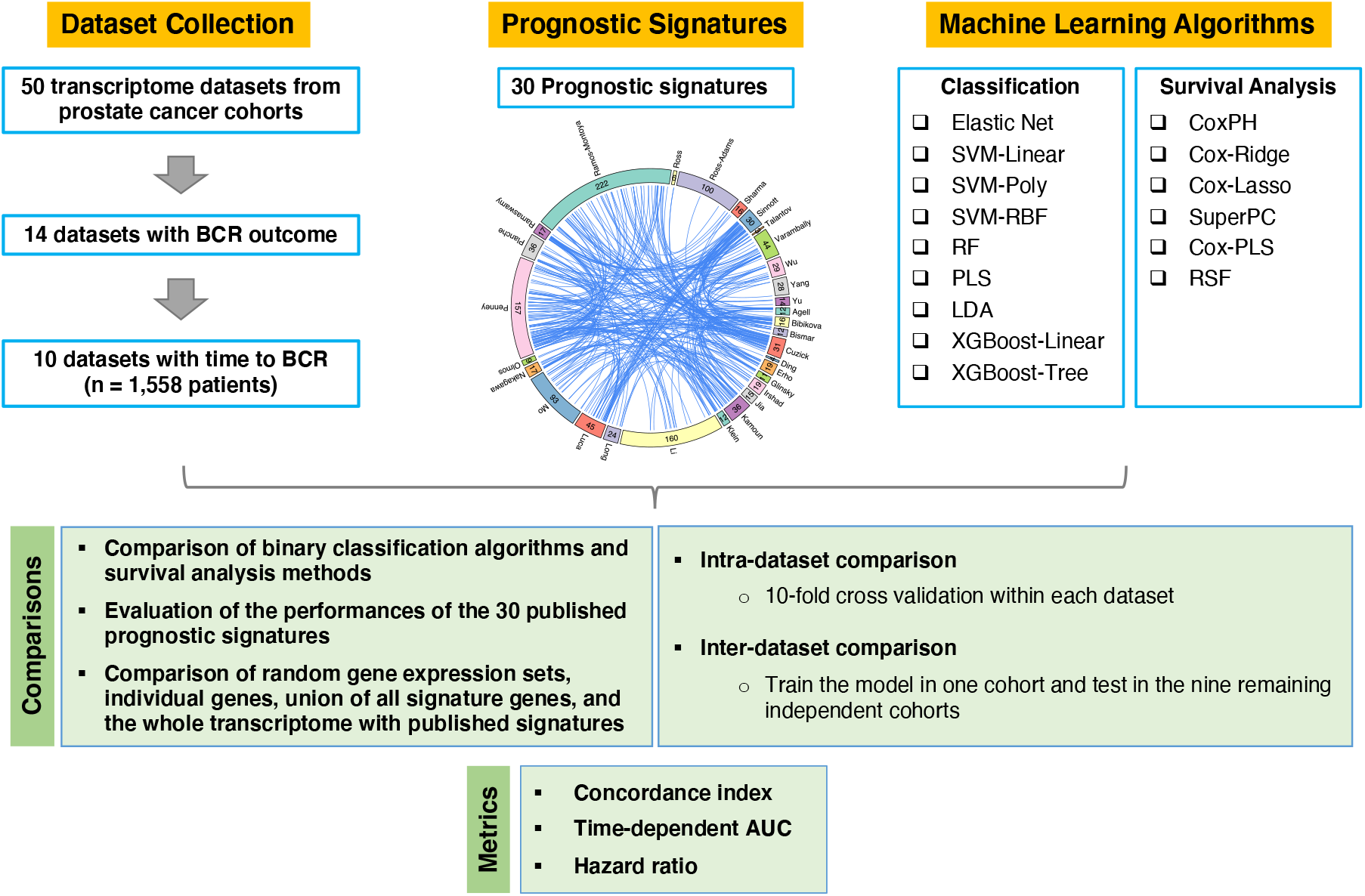
Overview of the study design. In general, the transcriptomics data and sample metadata for a total of 10 independent prostate cancer (PCa) cohorts (1,588 primary tumor samples) with the time to biochemical recurrence (BCR) data were collected and included in the study. The performances of 15 machine learning (ML) algorithms including 6 survival analysis methods and 9 binary classification algorithms were compared to identify the most robust one, and the 30 published gene expression signatures were analyzed and evaluated based on the most robust ML algorithm across all the independent cohorts. Both the intra-dataset (10-fold cross-validation within a dataset) and inter-dataset (train the model in one dataset and test the model in the remaining independent datasets) comparisons were conducted to evaluate the models and signatures.

### Functional characterization of the signature genes

In total, there were 1,032 unique genes in the 30 published prognostic signatures for PCa (union of the signature genes). Gene expression profiles were compared between the primary tumor samples versus the tumor-adjacent normal samples by leveraging the TCGA-PRAD data. The result showed that 223 among the 1,032 genes (21%) were differentially expressed with the absolute fold change (FC) > 2 and the false discovery rate (FDR) < 0.01 (Figure 2A). The CoxPH survival analysis based on the TCGA-PRAD data showed that 558 out of the 1,032 genes (54%) were significantly associated with RFS with P values < 0.05 (Figure 2B). The 1,032 genes were significantly enriched in many important cancer-associated pathways, including the cell cycle pathway, apoptosis pathway, p53 signaling pathway, PI3K-Akt signaling pathway, and prostate cancer pathway, etc. (Figure 2C), indicating the biological importance of these signature genes. The genes involved in those pathways were usually included in many signatures (Supplementary Figure S1).

**Figure 2.**
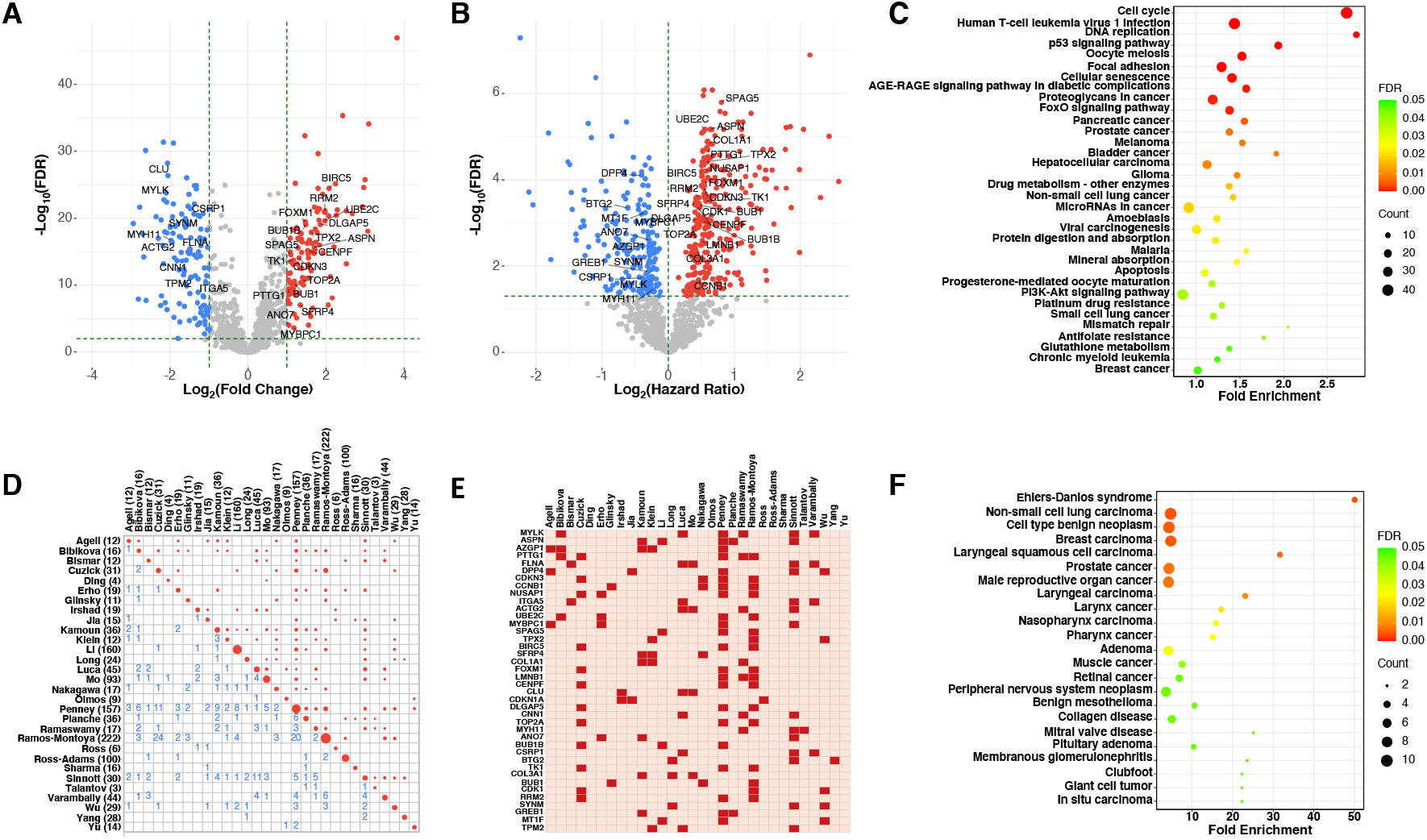
Functional characterization of the signature genes. (**A**) Differential expression analysis of the 1,042 unique signature genes (union of the 30 prognostic signatures) between tumor vs. normal samples based on the TCGA-PRAD data. Genes in more than 3 signatures are labeled. (**B**) Cox Proportional-Hazards (CoxPH) survival analysis of relapse-free survival (RFS) for the 1,042 genes based on the TCGA-PRAD data. Genes in more than 3 signatures are labeled. (**C**) Kyoto Encyclopedia of Genes and Genomes (KEGG) pathways that the 1,042 genes enriched in. **(D)** Overlap of genes between different prognostic signatures. (**E**) Heatmap showing the 40 common genes that were identified in three or more signatures. (**F**) Disease Ontology (DO) enrichment analysis of the 40 common genes identified in three or more signatures.

Analysis of the gene overlap among different signatures indicated that the prognostic signatures were largely distinct in terms of gene identity (Figure 2D). For example, a maximum of three genes were shared between the Erho signature (Decipher) and the Penney signature, while only one or two common genes were identified between the Erho signature and eight other signatures. Among the 1,032 genes, 142 genes (13.8%) were shared by more than two signatures and 40 genes (3.9%) were common in three or more signatures (Figure 2E). Based on the TCGA-PRAD data, 28 among the 40 common genes (70%) were significantly differentially expressed between the primary tumor samples versus the tumor-adjacent normal samples, and 36 out of the 40 common genes (90%) were significantly associated with RFS. The disease ontology analysis showed that these 40 genes were primarily associated with cancers, including prostate cancer (Figure 2F). Taken together, the results revealed that many the signature genes were indeed relevant to PCa and disease outcomes.

### Comparison of survival analysis models versus binary classification models

We first compared the performances of nine binary classification algorithms and six survival analysis methods in risk assessment. The sample size of each dataset for binary classification and survival analysis were summarized in Supplementary Table S5. For each algorithm, the 30 published signatures will be used individually to build the prediction model in the training dataset and then be validated in the validation datasets. For instance, in the intra-dataset comparison, we used 10-fold cross-validation to test a ML algorithm based on each of the 30 signatures across 10 datasets, whereas in the inter-dataset comparison, one dataset was used to train the 30 prognostic models and the remaining nine datasets were used for validations. In each strategy, the median values of the three metrics, i.e., C-index, AUC, and HR, were calculated across all the signatures and all the datasets to rank these ML algorithms. The results indicated that almost all the survival models were superior to binary classification models in the intra-dataset comparison (Figure 3), i.e., the median values of C-index, AUC, and HR across all the signatures and all the datasets for the survival models were greater than those for the binary classification models. The results in the inter-dataset comparison were generally consistent with those in the intra-dataset comparison. Some of the top binary classification models such as PLS, Elastic Net, and RF performed equally well or slightly better than the CoxPH or SuperPC survival models, however, the top survival models including Cox-Ridge, Cox-PLS, and RFS outperformed all the binary classification models in the inter-dataset comparison.

**Figure 3.**
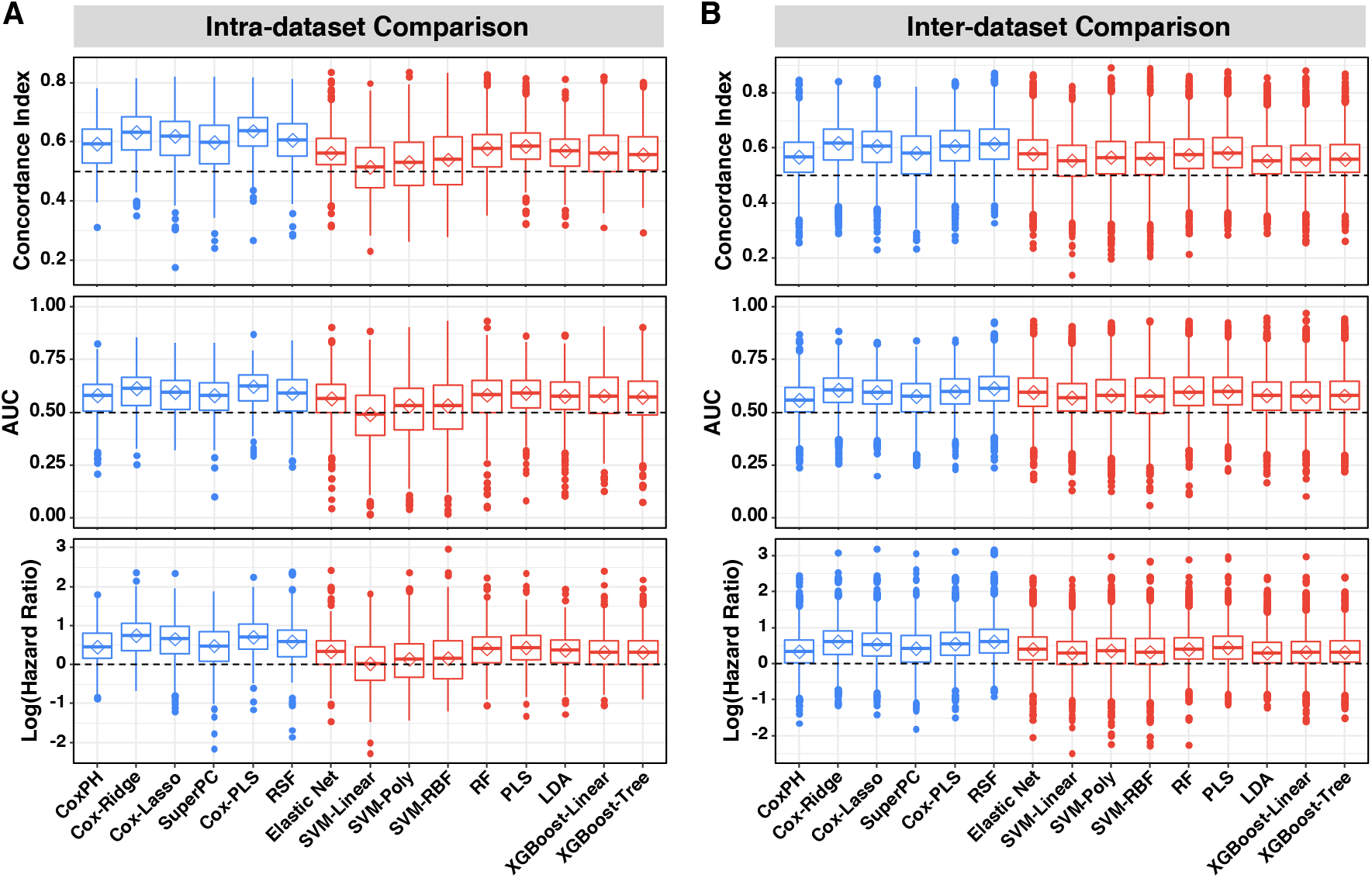
Comparison of survival analysis models versus binary classification models. Prognostic performances of the machine learning (ML) algorithms were evaluated based on all the 30 gene expression signatures across all the transcriptomics datasets. Both intra-dataset (**A**) and inter-dataset (**B**) comparisons were performed. Median values of the three metrics, including concordance index (C-index, *top*), area under the time-dependent receiver operating characteristic curve (AUC, *middle*), and hazard ratio (HR, *bottom*) estimated by the Kaplan Meier (KM) analysis of relapse-free survival (RFS) were used to rank the algorithms, respectively. The diamond in each box indicates the median value. Different colors of the box plots denote different categories of ML algorithms, *i*.*e*., blue for the six survival analysis methods and red for the nine binary classification algorithms.

There are two possible reasons why the survival analysis models were superior to the binary classification models: (i) the sample size became smaller in binary classification models if the censored data were excluded (Supplementary Table S5), and (ii) the variability in survival time provides more information than binary outcome, leading to increased data resolution and therefore statistical power. To test these hypotheses, we compare the survival models with the binary classification models using the same data for which the patients with censored data were removed. The results indicated that the prediction performances for all the survival models increased as sample size (Supplementary Figure S2), and the survival models also outperformed the binary classification models based on the same sample sizes (Supplementary Figure S3), especially in the intra-dataset comparison, which supported our hypotheses.

### Performance evaluation of the survival analysis methods

A comprehensive evaluation of the performances of the six algorithms for survival analysis were carried out (Figure 4). For the intra-dataset comparison based on 10-fold CV, Cox-Ridge and Cox-PLS models generally performed better than the other models, whereas the most widely used CoxPH method usually ranked bottom among six survival analysis algorithms. The performances of the SuperPC and RSF algorithms substantially varied and largely depended on the datasets that were used for analysis. For example, based on the C-index, the performances of RSF were very similar to the top 2 algorithms Cox-Ridge and Cox-PLS in the DKFZ, GSE54460, and Cambridge datasets, whereas the performances of RSF ranked the last or the second last in the CPC-Gene, Stockholm, CancerMap, CIT, and Belfast datasets. The performances of Cox-Lasso usually ranked higher than RSF, SuperPC, and CoxPH, but lower than Cox-Ridge and Cox-PLS. The result was slightly different for the inter-dataset comparison, where the performances of Cox-Ridge, Cox-PLS, and RSF were comparable and generally better than the other algorithms. The rankings of Cox-Lasso, CoxPH, and SuperPC were consistent with that in the intra-dataset comparison, where Cox-Lasso > SuperPC = CoxPH.

**Figure 4.**
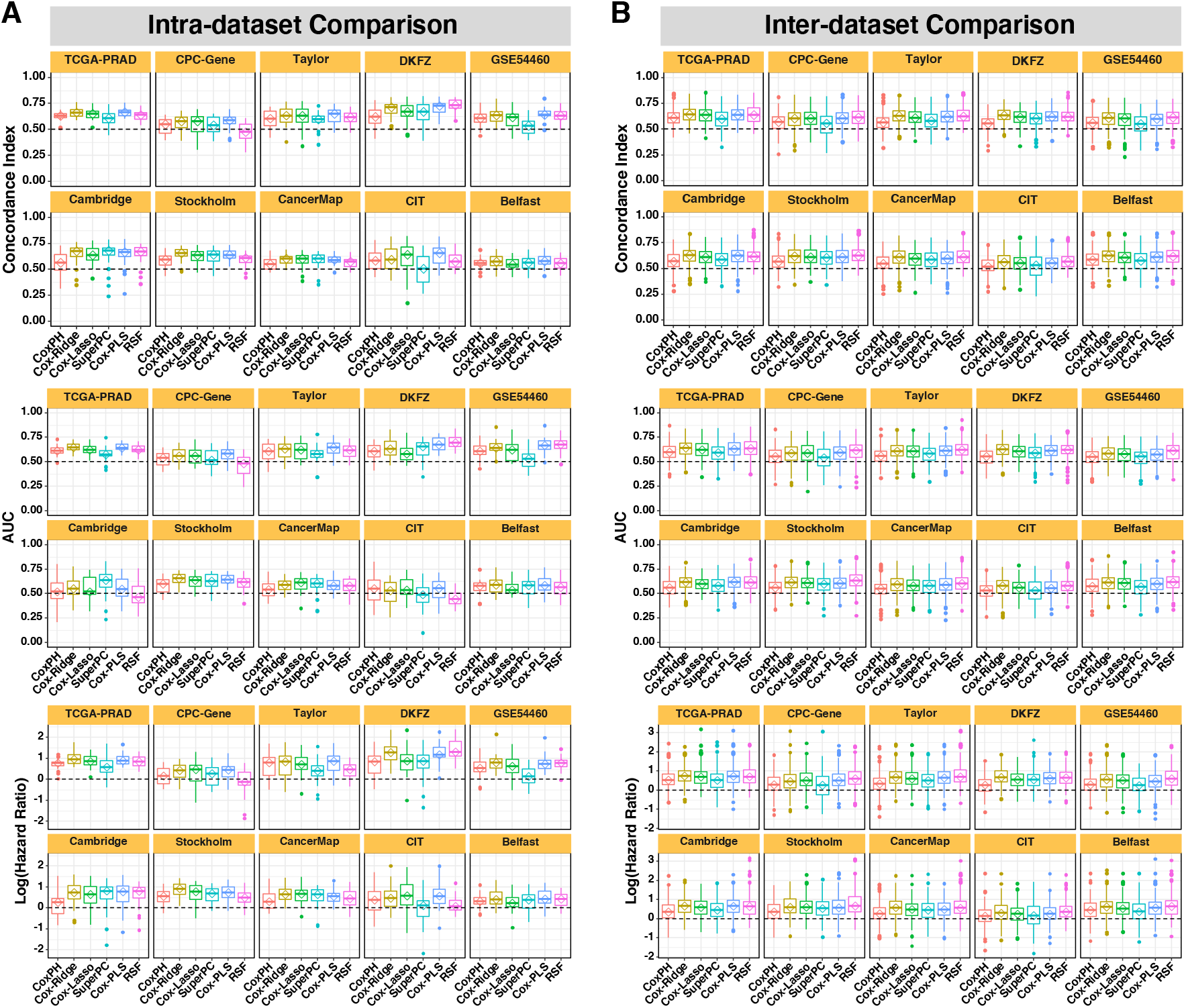
Comprehensive evaluation of the six survival analysis methods. The performances of the methods were evaluated based on all the 30 gene expression signatures within each individual dataset via 10-fold cross-validation for intra-dataset comparison (**A**), or using an individual dataset to train the prognostic models and the remaining independent datasets to test the models for inter-dataset comparison (**B**). The title of each small panel indicates the dataset (or the training dataset for inter-dataset comparison) that was being used for model comparison. Median values of the three metrics, including concordance index (C-index, *top*), area under the time-dependent receiver operating characteristic curve (AUC, *middle*), and hazard ratio (HR, *bottom*) estimated by the Kaplan Meier (KM) analysis of relapse-free survival (RFS) were used to rank the survival analysis methods, respectively. The diamond in each box indicates the median value.

It was as expected that the survival analysis models usually performed better in the intra-dataset comparison than that in the inter-dataset comparison. This was mainly because that in the intra-dataset comparison, the training and test data were generated in the same study using the same technology, whereas in the inter-dataset comparison, different technologies, different platforms, and even different bioinformatics pipelines may be used to generate the training and test sets. The cohorts may also be quite different between studies, yielding increased level of heterogeneity in inter-dataset analysis. These results in the study emphasized the importance of validation with multiple independent cohorts when building and evaluating prognostic models.

### Evaluation of the published gene expression signatures for PCa prognosis

Based on the comprehensive comparisons for different survival analysis algorithms, Cox-Ridge, which performed consistently well across all the datasets was selected to further evaluate the performances of various prognostic signatures. The median values of the three metrics were calculated across all the datasets and were used to rank the signatures. The results indicated that almost all the signatures had some prognostic powers with median C-indexes and median AUCs greater than 0.5, and median HRs greater than 1 both in the intra-dataset and in the inter-dataset comparisons (Figure 5). Some prognostic signatures, including Penny, Li, Klein, Sinnott, Wu, Erho, Kamoun, Planche, and Long, almost always ranked among the top 10 signatures based on the three metrics. Two of the three commercially applied prognostic signatures, i.e., Klein (Oncotype DX GPS) and Erho (Decipher), performed consistently well across the datasets, especially that Klein always ranked among the top 5 signatures in the intra-dataset comparison and ranked the second in the inter-dataset comparison. The median values of C-index, AUC, and HR for the signature Penney usually ranked the first, which was the only signature performing better than the Klein signature in most comparisons. Some of the other top signatures including Li, Sinnott, and Wu always had higher rankings than the Erho signature in the intra-dataset comparison, and Wu also outperformed the Erho signature in the inter-dataset comparison.

**Figure 5.**
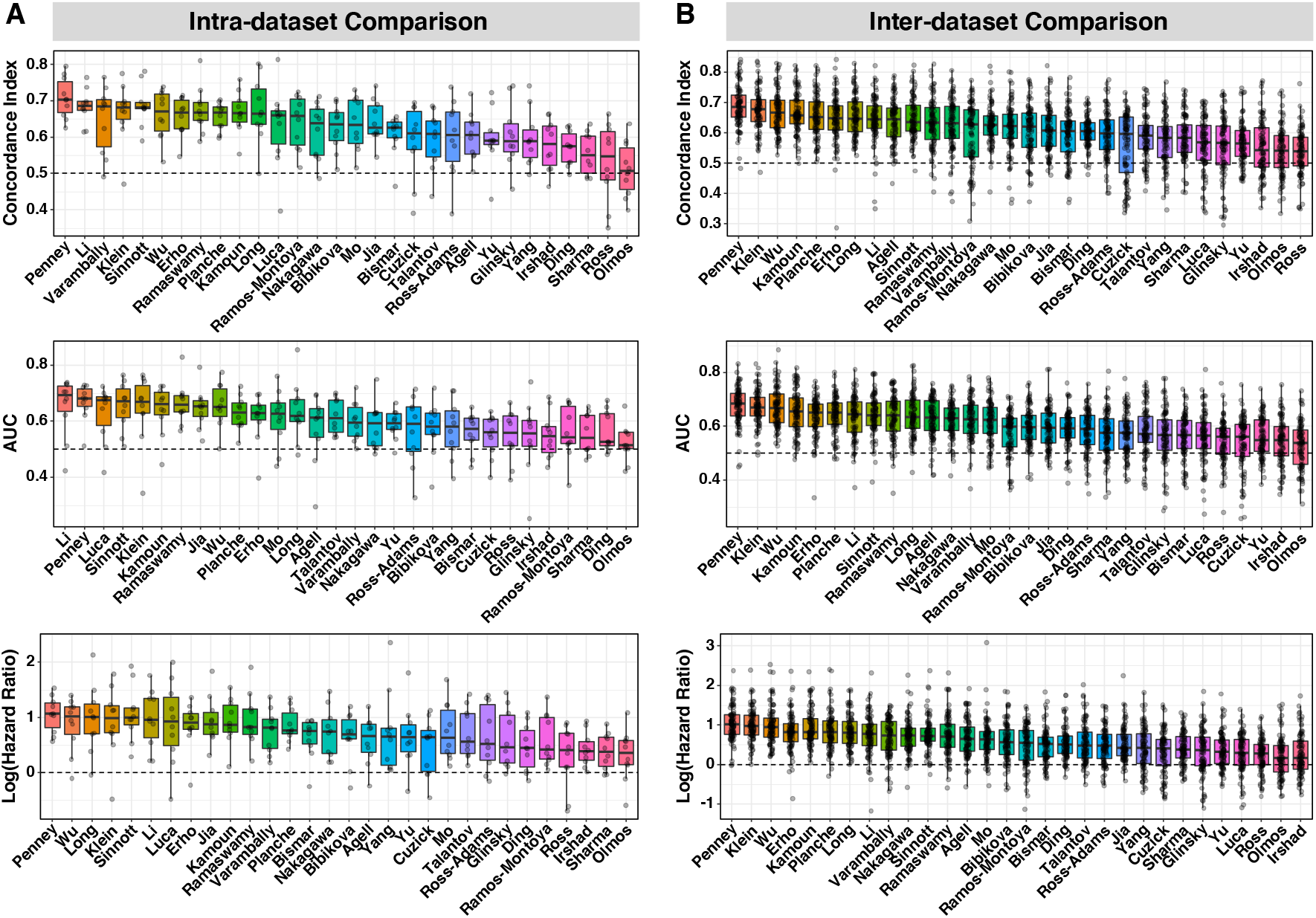
Evaluation of the 30 gene expression prognostic signatures based on the robust Cox-Ridge algorithm. (**A**) Intra-dataset comparison via 10-fold cross-validation. (**B**) Inter-dataset comparison using one dataset as the training set and the remaining datasets as test sets. The prognostic signatures are ranked based on the median values of the three metrics, including concordance index (C-index, *top*), area under the time-dependent receiver operating characteristic curve (AUC, *middle*), and hazard ratio (HR, *bottom*) estimated by the Kaplan Meier (KM) analysis of relapse-free survival (RFS), respectively, in the plots. Each dot represents the C-index, AUC, or HR value calculated within the dataset via 10-fold cross-validation for the intra-dataset comparison or in a test dataset for the inter-dataset comparison.

We also did similar comparisons using the other well-performed algorithm Cox-PLS to investigate whether the performances of the signatures relied on the methods for model development. The results indicated that although the rankings may be slightly different based on different algorithms, the top signatures using the Cox-Ridge algorithm also ranked in the top list when Cox-PLS was used (Supplementary Figure S4). The signature Ramaswamy performed slightly better using the Cox-PLS algorithm, which made it always ranked among the top 10 signatures.

Since the test characteristics, i.e., C-index, AUC, and HR, may vary when different datasets were used as training and test sets, we utilized the rankings of the median values of these three metrics across all possible combinations of training and test sets to reflect the overall performances of the signatures being evaluated. Figure 6 showed details about the HR-based performances of the signatures for each dataset in the intra-dataset comparison, while Figure 7 revealed such performances when each dataset was used as training set in the inter-dataset comparison. The rankings of the top signatures such as Penny, Li, Klein, Sinnott, Wu, Erho, etc. were generally high across multiple comparisons. For instance, in the intra-dataset comparison, the signature Sinnott always ranked among the top five signatures in six datasets, including CPC-Gene, Taylor, DKFZ, GSE54460, CancerMap, and Belfast, whereas the signature Penney ranked among the top five signatures in five datasets, including GSE54460, Cambridge, Stockholm, CancerMap, and Belfast (Figure 6). In the inter-dataset comparison, the signature Penney and Klein can be independently validated in more than half of the nine test datasets no matter which training dataset was used when building the model (Figure 7). The signature Wu can also be validated by multiple independent cohorts when different training sets were used. Supplementary Figure S5 showed the examples of survival curves for the top signatures - Penney and Wu, and the two commercial panels – Klein and Erho when the TCGA-PRAD dataset was used as the training set and the other datasets as validation sets. We also observed that some signatures may perform well when certain datasets were used for model development and validation, but they could show very low or even no prognostic power when other datasets were tried. The results suggested the importance of leveraging data from multiple cohorts to develop and validate new prognostic models or to evaluate existing prognostic models.

**Figure 6.**
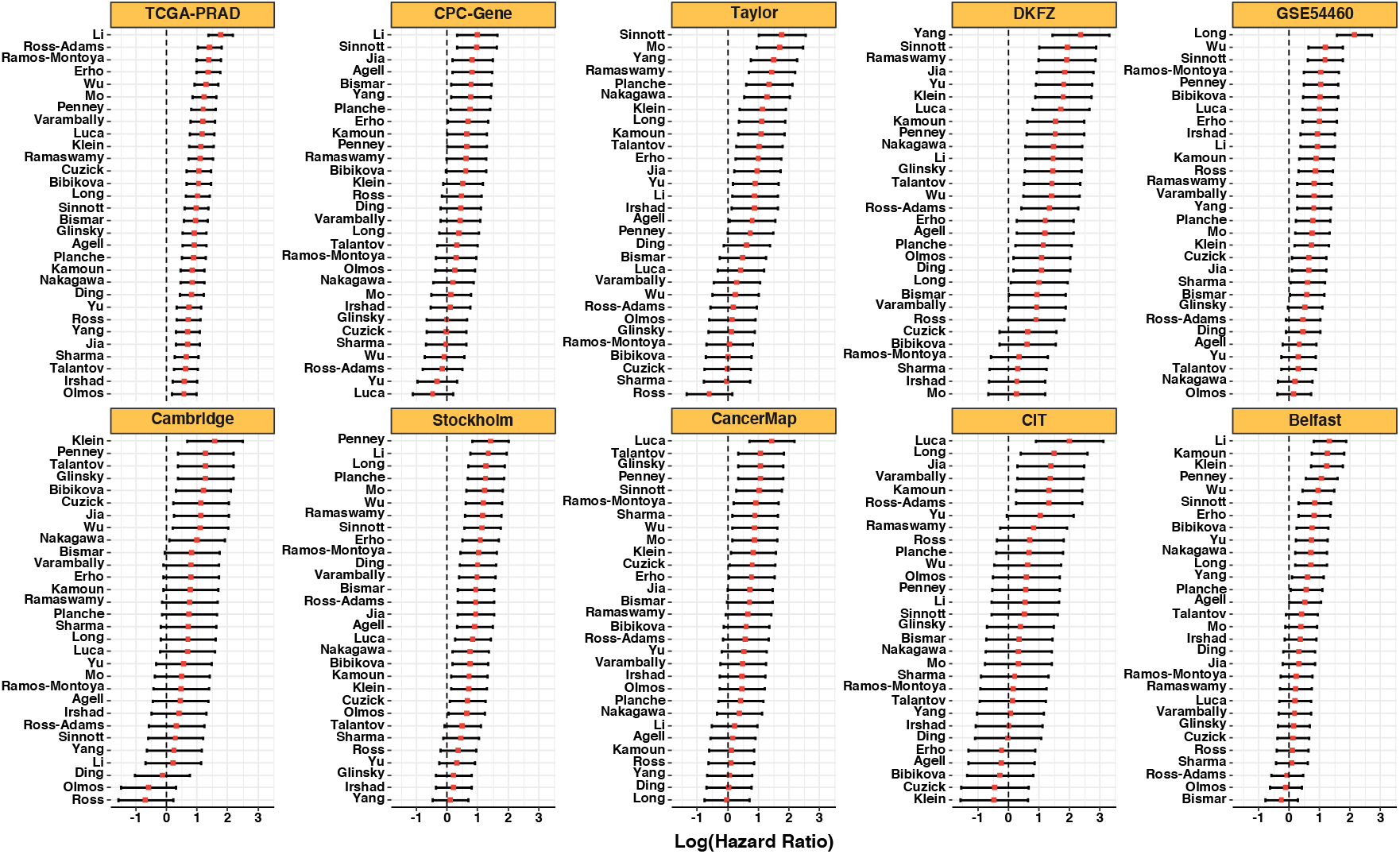
Intra-dataset comparison of the 30 gene expression prognostic signatures based on hazard ratio (HR) estimated by the Kaplan Meier (KM) analysis of relapse-free survival (RFS). The robust Cox-Ridge algorithm was used to train the prognostic models. The signatures are ranked by HRs in each individual dataset in the plots. The title of each small panel indicates the name of the dataset.

**Figure 7.**
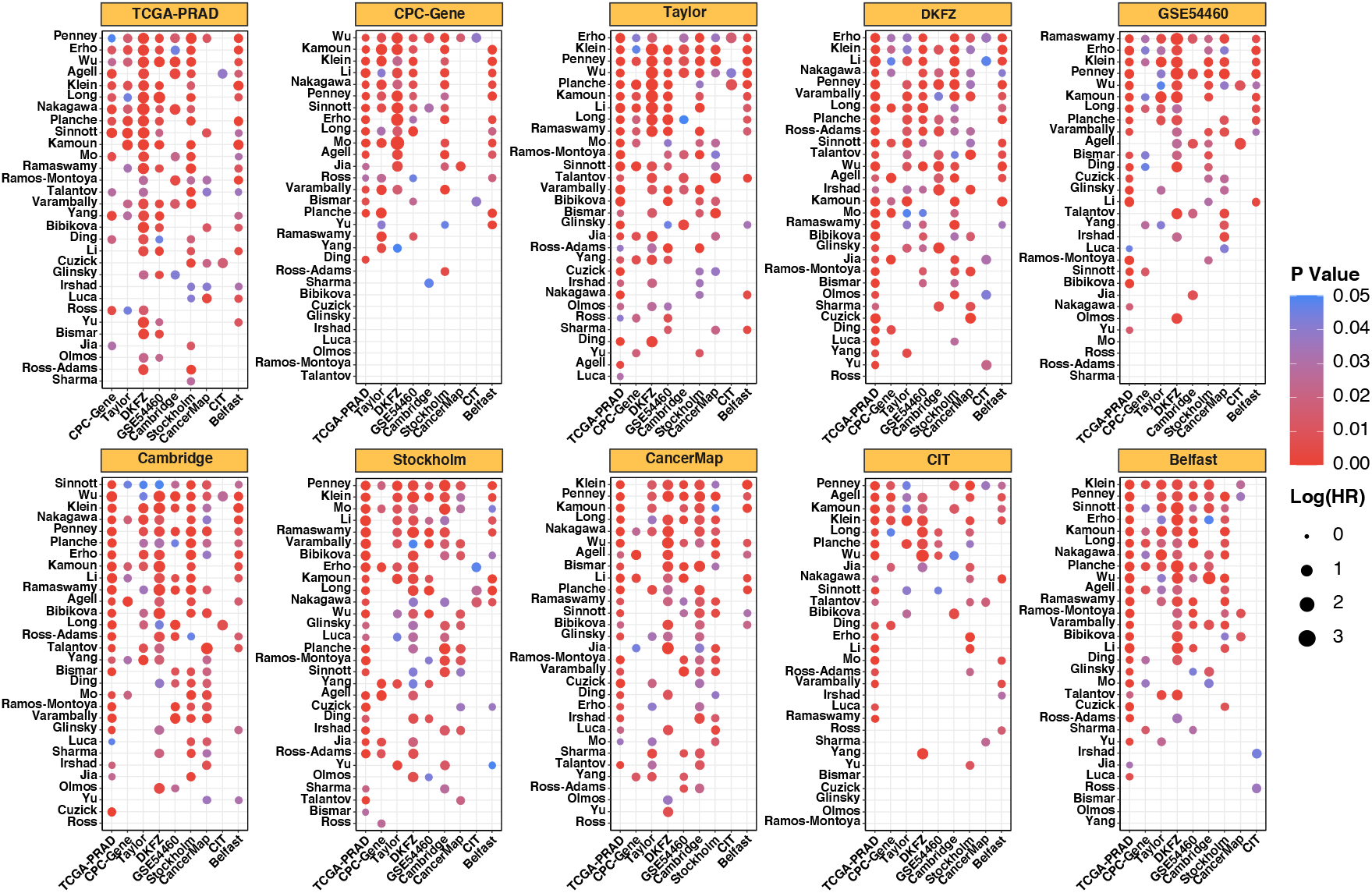
Inter-dataset comparison of the 30 gene expression prognostic signatures based on hazard ratio (HR) estimated by the Kaplan Meier (KM) analysis of relapse-free survival (RFS). The robust Cox-Ridge algorithm was used to train the prognostic models. The title of each small panel refers to the training dataset. The x-axis indicates the test datasets and the y-axis indicates the signatures. The prognostic signatures are ranked based on the numbers of test datasets in which the prognostic models have been successfully validated (p value < 0.05) based on KM survival analysis. Only the validations with p value < 0.05 are shown in the figure.

### The necessity and importance of deriving molecular signatures from gene expression profiling

We compared the performances of the published signatures with RGESs, the whole transcriptome, individual signature genes, and union of genes in the published signatures to investigate the necessity of deriving gene expression signatures for PCa prognosis. We generated 10 RGESs with 50 genes each and performed both the intra-dataset and inter-dataset analyses as what have been done for the published signatures using the Cox-Ridge algorithm. Consistent with some previous studies (16,17), RGESs could be significantly associated with clinical outcomes of cancer, however, it was observed that the top published signatures always outperformed all the 10 RGESs in our study (Supplementary Figure S6). We also investigated the performances of prediction models leveraging the whole transcriptome without any feature selection for PCa prognosis. The results showed that the prognostic models using the whole transcriptome generally performed reasonably well, which were better than two third of the published signatures, but they were not as good as the top signatures (Supplementary Figure S7). To investigate whether the individual genes in a published signature have comparable performances with the entire gene expression signature in risk assessment, the expression values of each individual gene were used as the risk scores and the C index, AUC, and HR of KM analysis were calculated in each dataset. These metrics for individual genes were compared with those for the published signatures calculated in the intra-dataset comparison using the robust Cox-Ridge algorithm. As expected, the top 10 individual signature genes performed better than some of the published signatures, however, none of them outperformed the top signatures (Supplementary Figure S8). The union of the signature genes was also modeled using the same strategy and compared with the published signatures to test if integrated gene sets lead to a better performance. The results showed that the performances of the signature with union genes were comparable with the top signatures, which always ranked in the top 5 across all the comparisons (Supplementary Figure S9). On the other hand, it also indicated that a relatively small set of the most significant genes is able to achieve as high prognostic accuracy as the large signatures do, and will be much easier to be translated into clinical practice when the analytical validity and cost-effectiveness are taken into consideration. The sum of the evidences indicated that it is desirable to develop PCa prognostic models using the most significant outcome-associated genes selected from transcriptome.

### Translational potential of the prognostic signatures in multiple race groups

Although most published prognostic signatures for PCa were developed and validated in cohorts of predominantly white men, the signatures could also be significantly associated with clinical outcomes of PCa patients in other race/ethnicity populations, thus may potentially be validated in well-designed clinical trials to assist on clinical decision-making for non-white men with PCa. To test this hypothesis, we carried out an analysis by leveraging the TCGA-PRAD dataset that consists of primary PCa samples from 386 white men, 52 African American men and 12 Asian men. The robust Cox-Ridge algorithm was used to develop prediction models based on each of the 30 signatures using the 386 training samples of white men. The models were then validated by the cohort of African American patients only (N=52) and by the combined cohort of the African American patients and the Asian patients (N=64), respectively. KM survival analysis was performed to evaluate the associations of the 30 published signatures with the time to BCR. The results indicated that 12 signatures can be validated in the African American cohort and 19 signatures can be validated in the combined cohort (African Americans + Asians) with p values less than 0.05, including many of the top signatures in the aforementioned study such as Li, Penney, Wu, Erho, and Klein, as well as some other signatures such as Ross-Adams, Romos-Montoya, and Varambally, etc. (Supplementary Figure S10). In spite of small sample size, the promising results warrant further assessment of these signatures with multiple large validation cohorts of different races and ethnicities.

## Discussion

Unlike many other cancer types, a major challenge of localized PCa management is to accurately predict the risk of the disease aggressiveness at the time of diagnosis to make informed clinical decisions to avoid over-treatment because the majority of localized PCa tumors grow slowly and will likely never cause health problems but many patients will still undergo RP, a surgical procedure to remove the entire prostate gland, which may adversely impact health-related quality of life. Accurate risk assessment tools are desperately needed to assist on PCa treatment decision-making. Molecular signatures derived from gene expression profiling are becoming critical tools and have impacted the clinical practice in PCa management. Once validated in various clinical settings, the molecular signatures will be of great utility, including (i) identification of candidates for active surveillance to avoid unnecessary surgery and selection of patients who will need immediate treatment to avoid missing the opportunity for early intervention. (ii) determination of the type and intensity of treatment, (iii) evaluation of the need of multimodal therapy or the possible benefits of adjuvant therapies, and (iv) periodic monitoring of local recurrence and distant metastasis after definitive therapy.

A systematic and unbiased evaluation of ML algorithms and gene expression signatures for cancer prognosis using large cohorts is very critical to select the optimal approaches for building the predictive models and identify the most promising signatures for further validations in prospective clinical studies before they can be applied to clinical practice. The results from our study revealed that the survival analysis algorithms modeling the censored time-to-event data generally outperformed the binary classification algorithms. This is mainly because that the censored data were removed when building the binary classifiers which resulted in smaller sample size, and the binary classification algorithms themselves were not as good as the survival analysis algorithms for risk assessment. Among the six survival analysis methods, Cox-Ridge and Cox-PLS generally performed better than the other algorithms, whereas the most widely used method CoxPH usually performed the worst or the second worst. The performances of RSF were not good in the intra-dataset comparison, but were very robust in the inter-dataset comparison. It was as expected that the prediction models in the inter-dataset comparison usually do not perform as well as that in the intra-dataset comparison because of the high heterogeneities between different cohorts. This emphasized the importance of clinical validations in independent cohorts for the development and evaluation of prognostic models.

In this study, the two commercial tests, Oncotype DX GPS (Klein signature) and Decipher (Erho signature) almost always ranked in the top signatures in all the comparisons. We were also able to identify a few other published signatures such as Penney, Wu, Li, and Sinnott that generally had comparable or even better performances than the commercial panels. These four signatures and some other top signatures such as Kamoun, Planche, and Long could be promising for developing robust prognostic models to be incorporated into clinical practice with further prospective validations. The promising signatures were identified mainly based on the overall performances. It’s possible that a signature could perform well using a given machine learning algorithm and a given training data to develop the predictive model, while could have no prognostic power when other algorithms or training datasets were used. Therefore, multiple robust survival analysis algorithms and large PCa population cohorts were recommended when developing new prognostic signatures. A few previous studies have demonstrated that RGESs may be significantly associated with clinical outcomes of cancer, which has also been observed in our study, however, the top published signatures always outperformed all RGESs. We also evaluated the performances of prognostic models using the whole transcriptome data and found that the transcriptome models could outperform some of the signatures but were not comparable with the top signatures. Although the performance of the union of the signature genes were comparable with the top signatures and in certain cases were slightly better than them, it would be challenging to translate the signatures with a large number genes into clinical practice. These findings from our study indicated that it’s critical to derive molecular signatures with a relatively small number of genes that are relevant to PCa outcomes from the transcriptome to further improve the PCa prognosis to assist on clinical decision-making.

From the comprehensive review and evaluation of the published prognostic signatures for PCa, we proposed a few practice principles should be followed when developing and evaluating a molecular signature for PCa prognosis, such as (i) demonstration of clinical utility, (ii) selection of the most relevant cohorts and endpoints, (iii) selection of the robust machine learning algorithm and modeling time-to-event data if available rather than categorical data, (iv) conducting multiple external validations to demonstrate the clinical validity, and (v) comparison with the top published signatures and the established clinico-pathological prognostic markers. Some of these principles have also been extensively reviewed and discussed elsewhere (47–49). Clinical utility is the most important criterion for molecular signature development. The majority of the published gene expression signatures for PCa prognosis have been developed based on RP tissue in surgical cohorts. Thus, these signatures may be more relevant to patients with primary treatment for assessing the risk of recurrence, metastasis, or prostate cancer– specific mortality (PCSM), and determining if the patients will benefit from adjuvant therapy. Although some of these signatures have also been proven to be useful to identify candidates for active surveillance with prostate needle biopsy samples, the signatures directly derived in the prostate needle biopsy setting are needed to boost sensitivity and specificity of the test. It’s also critical to select the most relevant endpoints for future molecular signature development and validation. Some published signatures were developed to predict the Gleason score, which may have limited clinical impact. The surrogate endpoint BCR is most commonly used to define disease progression or aggressiveness, however, it should be noted that many patients with an increase of PSA level following the definitive therapy may not experience local recurrence or develop distant metastasis for a few years. The clinical recurrence, metastasis, and PCSM may be better endpoints, and if possible, the time-to-event data should be modeled using the most robust survival analysis algorithms to improve the accuracy of risk stratification. The endpoint adverse pathology could also be used to define the disease aggressiveness when developing molecular signatures for risk assessment at the time of biopsy. Clinical validity is another critical criterion for molecular signature development. Prognostic signatures are of no utility without multiple well-designed external validations. The signature scores computed using any models will be significantly associated with the endpoint of interest in the training dataset. The findings from our study emphasized the importance of independent validation when building and evaluating prognostic models. In addition, any new prognostic signatures should be compared with the top published signatures and the established clinico-pathological prognostic markers to assess the incremental values of the signatures.

In summary, this is the first study that performed a comprehensive evaluation of the performances of multiple machine learning models and published gene expression signatures using large PCa cohorts. The findings from the study have greatly facilitated the identification of existing prognostic signatures for further validations and the selection of the best strategies for the development of more robust signatures to assist on PCa treatment decision making. In addition, our study also provided a valuable resource with 10 transcriptomics datasets from large primary PCa cohorts and a comprehensive collection of 30 gene expression prognostic signatures that can be used to develop, validate, and evaluate novel statistical methodologies and molecular signatures to improve PCa management.

## Availability of data and materials

The processed data including the normalized expression data and the harmonized metadata of all the 10 transcriptomics datasets were deposited in the PCaDB database (http://bioinfo.jialab-ucr.org/PCaDB/) and the *ExpressionSet* object of each dataset can be easily downloaded from this database. All the scripts used in this study, including data preprocessing, model comparison, and data visualization, are publicly available at https://github.com/rli012/PCaSignatures.

## References

1. Sung H, Ferlay J, Siegel RL, Laversanne M, Soerjomataram I, Jemal A, et al. Global Cancer Statistics 2020: GLOBOCAN Estimates of Incidence and Mortality Worldwide for 36 Cancers in 185 Countries. CA: A Cancer Journal for Clinicians. 2021;71:209–49.

2. Hruza M, Bermejo JL, Flinspach B, Schulze M, Teber D, Rumpelt HJ, et al. Long-term oncological outcomes after laparoscopic radical prostatectomy. BJU International. 2013;111:271–80.

3. Freedland SJ, Humphreys EB, Mangold LA, Eisenberger M, Dorey FJ, Walsh PC, et al. Risk of Prostate Cancer–Specific Mortality Following Biochemical Recurrence After Radical Prostatectomy. JAMA. 2005;294:433–9.

4. Stephenson AJ, Kattan MW, Eastham JA, Dotan ZA, Bianco FJ, Lilja H, et al. Defining biochemical recurrence of prostate cancer after radical prostatectomy: a proposal for a standardized definition. J Clin Oncol. 2006;24:3973–8.

5. Bhargava P, Ravizzini G, Chapin BF, Kundra V. Imaging Biochemical Recurrence After Prostatectomy: Where Are We Headed? American Journal of Roentgenology. American Roentgen Ray Society; 2020;214:1248–58.

6. Erho N, Crisan A, Vergara IA, Mitra AP, Ghadessi M, Buerki C, et al. Discovery and validation of a prostate cancer genomic classifier that predicts early metastasis following radical prostatectomy. PLoS One. 2013;8:e66855.

7. Irshad S, Bansal M, Castillo-Martin M, Zheng T, Aytes A, Wenske S, et al. A Molecular Signature Predictive of Indolent Prostate Cancer. Science Translational Medicine. American Association for the Advancement of Science; 2013;5:202ra122–202ra122.

8. Cuzick J, Swanson GP, Fisher G, Brothman AR, Berney DM, Reid JE, et al. Prognostic value of an RNA expression signature derived from cell cycle proliferation genes in patients with prostate cancer: a retrospective study. Lancet Oncol. 2011;12:245–55.

9. Long Q, Xu J, Osunkoya AO, Sannigrahi S, Johnson BA, Zhou W, et al. Global transcriptome analysis of formalin-fixed prostate cancer specimens identifies biomarkers of disease recurrence. Cancer Res. 2014;74:3228–37.

10. Wu C-L, Schroeder BE, Ma X-J, Cutie CJ, Wu S, Salunga R, et al. Development and validation of a 32-gene prognostic index for prostate cancer progression. Proc Natl Acad Sci U S A. 2013;110:6121–6.

11. Luca B-A, Brewer DS, Edwards DR, Edwards S, Whitaker HC, Merson S, et al. DESNT: A Poor Prognosis Category of Human Prostate Cancer. Eur Urol Focus. 2018;4:842–50.

12. Ross-Adams H, Lamb AD, Dunning MJ, Halim S, Lindberg J, Massie CM, et al. Integration of copy number and transcriptomics provides risk stratification in prostate cancer: A discovery and validation cohort study. EBioMedicine. 2015;2:1133–44.

13. You S, Knudsen BS, Erho N, Alshalalfa M, Takhar M, Al-Deen Ashab H, et al. Integrated Classification of Prostate Cancer Reveals a Novel Luminal Subtype with Poor Outcome. Cancer Res. 2016;76:4948–58.

14. Klein EA, Cooperberg MR, Magi-Galluzzi C, Simko JP, Falzarano SM, Maddala T, et al. A 17-gene assay to predict prostate cancer aggressiveness in the context of Gleason grade heterogeneity, tumor multifocality, and biopsy undersampling. Eur Urol. 2014;66:550–60.

15. Zhao X, Rødland EA, Sørlie T, Vollan HKM, Russnes HG, Kristensen VN, et al. Systematic assessment of prognostic gene signatures for breast cancer shows distinct influence of time and ER status. BMC Cancer. 2014;14:211.

16. Venet D, Dumont JE, Detours V. Most Random Gene Expression Signatures Are Significantly Associated with Breast Cancer Outcome. PLOS Computational Biology. Public Library of Science; 2011;7:e1002240.

17. Tang H, Wang S, Xiao G, Schiller J, Papadimitrakopoulou V, Minna J, et al. Comprehensive evaluation of published gene expression prognostic signatures for biomarker-based lung cancer clinical studies. Annals of Oncology. Elsevier; 2017;28:733–40.

18. Penney KL, Sinnott JA, Fall K, Pawitan Y, Hoshida Y, Kraft P, et al. mRNA Expression Signature of Gleason Grade Predicts Lethal Prostate Cancer. JCO. Wolters Kluwer; 2011;29:2391–6.

19. Li R, Wang S, Cui Y, Qu H, Chater JM, Zhang L, et al. Extended application of genomic selection to screen multiomics data for prognostic signatures of prostate cancer. Brief Bioinform. 2021;22:bbaa197.

20. Sinnott JA, Peisch SF, Tyekucheva S, Gerke T, Lis R, Rider JR, et al. Prognostic Utility of a New mRNA Expression Signature of Gleason Score. Clin Cancer Res. 2017;23:81–7.

21. Jensen MA, Ferretti V, Grossman RL, Staudt LM. The NCI Genomic Data Commons as an engine for precision medicine. Blood. 2017;130:453–9.

22. Gao J, Aksoy BA, Dogrusoz U, Dresdner G, Gross B, Sumer SO, et al. Integrative Analysis of Complex Cancer Genomics and Clinical Profiles Using the cBioPortal. Sci Signal. 2013;6:pl1.

23. Barrett T, Wilhite SE, Ledoux P, Evangelista C, Kim IF, Tomashevsky M, et al. NCBI GEO: archive for functional genomics data sets—update. Nucleic Acids Research. 2013;41:D991–5.

24. Brazma A, Parkinson H, Sarkans U, Shojatalab M, Vilo J, Abeygunawardena N, et al. ArrayExpress—a public repository for microarray gene expression data at the EBI. Nucleic Acids Research. 2003;31:68–71.

25. Li R, Qu H, Wang S, Wei J, Zhang L, Ma R, et al. GDCRNATools: an R/Bioconductor package for integrative analysis of lncRNA, miRNA and mRNA data in GDC. Bioinformatics. 2018;34:2515–7.

26. Carvalho BS, Irizarry RA. A framework for oligonucleotide microarray preprocessing. Bioinformatics. 2010;26:2363–7.

27. Dobin A, Davis CA, Schlesinger F, Drenkow J, Zaleski C, Jha S, et al. STAR: ultrafast universal RNA-seq aligner. Bioinformatics. 2013;29:15–21.

28. Liao Y, Smyth GK, Shi W. featureCounts: an efficient general purpose program for assigning sequence reads to genomic features. Bioinformatics. 2014;30:923–30.

29. Robinson MD, McCarthy DJ, Smyth GK. edgeR: a Bioconductor package for differential expression analysis of digital gene expression data. Bioinformatics. 2010;26:139–40.

30. Davis S, Meltzer PS. GEOquery: a bridge between the Gene Expression Omnibus (GEO) and BioConductor. Bioinformatics. 2007;23:1846–7.

31. Li R, Zhu J, Zhong W-D, Jia Z. PCaDB - a comprehensive and interactive database for transcriptomes from prostate cancer population cohorts. bioRxiv. Cold Spring Harbor Laboratory; 2021;2021.06.29.449134.

32. Luca B-A, Moulton V, Ellis C, Connell SP, Brewer DS, Cooper CS. Convergence of Prognostic Gene Signatures Suggests Underlying Mechanisms of Human Prostate Cancer Progression. Genes (Basel). 2020;11:E802.

33. Mo F, Lin D, Takhar M, Ramnarine VR, Dong X, Bell RH, et al. Stromal Gene Expression is Predictive for Metastatic Primary Prostate Cancer. Eur Urol. 2018;73:524–32.

34. Yang L, Roberts D, Takhar M, Erho N, Bibby BAS, Thiruthaneeswaran N, et al. Development and Validation of a 28-gene Hypoxia-related Prognostic Signature for Localized Prostate Cancer. EBioMedicine. 2018;31:182–9.

35. Yates AD, Achuthan P, Akanni W, Allen J, Allen J, Alvarez-Jarreta J, et al. Ensembl 2020. Nucleic Acids Research. 2020;48:D682–8.

36. Tweedie S, Braschi B, Gray K, Jones TEM, Seal RL, Yates B, et al. Genenames.org: the HGNC and VGNC resources in 2021. Nucleic Acids Research. 2021;49:D939–46.

37. Maglott D, Ostell J, Pruitt KD, Tatusova T. Entrez Gene: gene-centered information at NCBI. Nucleic Acids Research. 2005;33:D54–8.

38. Kuhn M. Building Predictive Models in R Using the caret Package. Journal of Statistical Software. 2008;28:1–26.

39. Simon N, Friedman J, Hastie T, Tibshirani R. Regularization Paths for Cox’s Proportional Hazards Model via Coordinate Descent. J Stat Softw. 2011;39:1– 13.

40. Bair E, Tibshirani R. Semi-Supervised Methods to Predict Patient Survival from Gene Expression Data. PLoS Biol. 2004;2:e108.

41. Bastien P, Bertrand F, Meyer N, Maumy-Bertrand M. Deviance residuals-based sparse PLS and sparse kernel PLS regression for censored data. Bioinformatics. 2015;31:397–404.

42. Ishwaran H, Kogalur UB, Blackstone EH, Lauer MS. Random survival forests. The Annals of Applied Statistics. Institute of Mathematical Statistics; 2008;2:841–60.

43. Schröder MS, Culhane AC, Quackenbush J, Haibe-Kains B. survcomp: an R/Bioconductor package for performance assessment and comparison of survival models. Bioinformatics. 2011;27:3206–8.

44. Heagerty PJ, Lumley T, Pepe MS. Time-dependent ROC curves for censored survival data and a diagnostic marker. Biometrics. 2000;56:337–44.

45. Ritchie ME, Phipson B, Wu D, Hu Y, Law CW, Shi W, et al. limma powers differential expression analyses for RNA-sequencing and microarray studies. Nucleic Acids Research. 2015;43:e47–e47.

46. Wu T, Hu E, Xu S, Chen M, Guo P, Dai Z, et al. clusterProfiler 4.0: A universal enrichment tool for interpreting omics data. The Innovation. 2021;100141.

47. Michiels S, Ternès N, Rotolo F. Statistical controversies in clinical research: prognostic gene signatures are not (yet) useful in clinical practice. Annals of Oncology. 2016;27:2160–7.

48. Teutsch SM, Bradley LA, Palomaki GE, Haddow JE, Piper M, Calonge N, et al. The Evaluation of Genomic Applications in Practice and Prevention (EGAPP) initiative: methods of the EGAPP Working Group. Genet Med. 2009;11:3–14.

49. Simon RM, Paik S, Hayes DF. Use of Archived Specimens in Evaluation of Prognostic and Predictive Biomarkers. JNCI: Journal of the National Cancer Institute. 2009;101:1446–52.

