## Supplementary Materials for "Comprehensive evaluation of machine learning models and gene expression signatures for prostate cancer prognosis using large population cohorts"

### Supplementary Figures

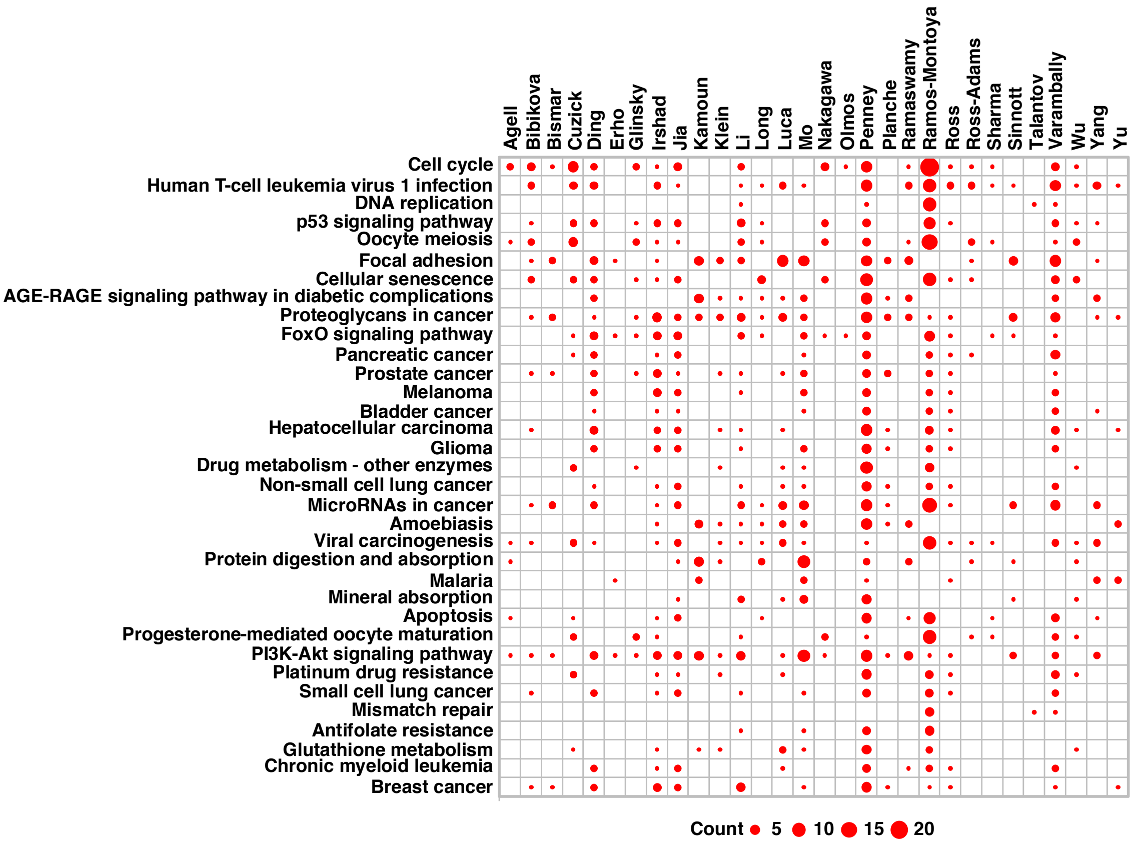


**Supplementary Figure S1.** The number of genes in each signature that are involved in the Kyoto Encyclopedia of Genes and Genomes（KEGG) pathways.

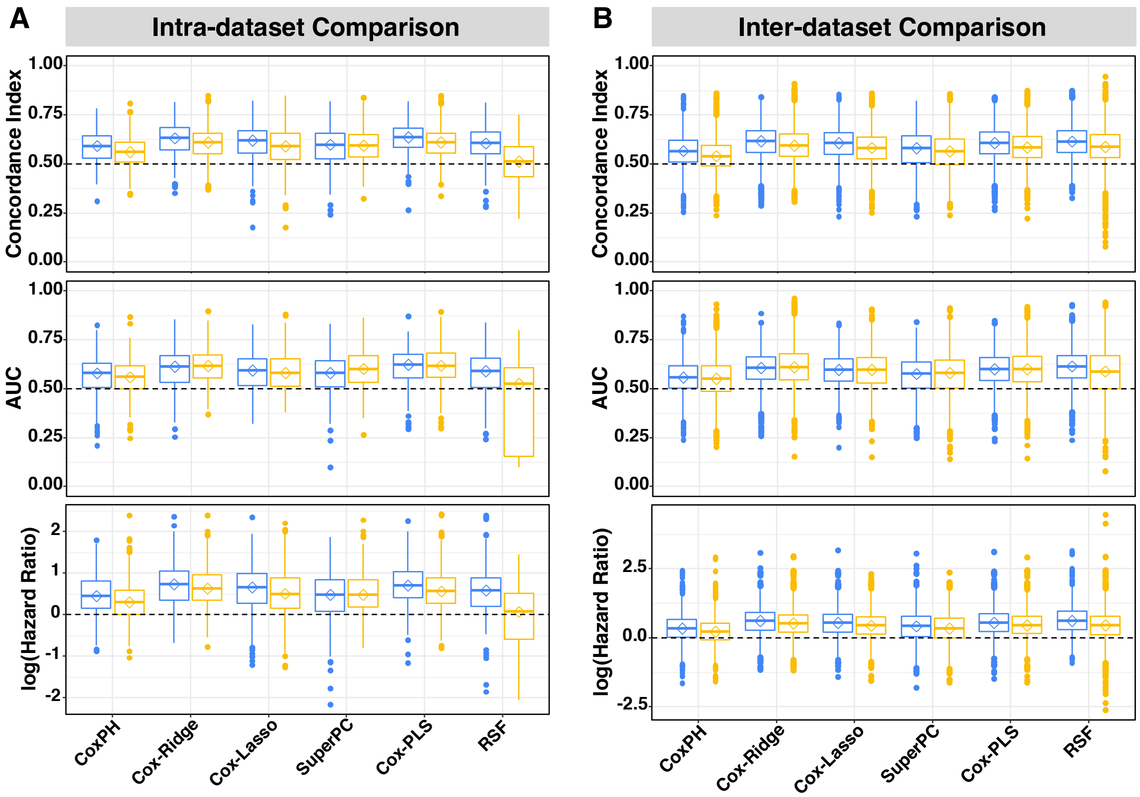


**Supplementary Figure S2.** Comparison of the six survival analysis models based on different sample sizes. Prognostic performances of the algorithms were evaluated across all the 30 gene expression signatures and across all the transcriptomics datasets. Both intra-dataset (**A**) and inter-dataset (**B**) comparisons were performed. Different colors of the box plots denote the analyses using different sample sizes, *i.e.*, blue for larger sample sizes with all the samples included and yellow for smaller sample sizes with the censored data (no BCR occurred within 5 years following primary treatment) removed. Median values of the three metrics, including Concordance index (C-index, *top*), area under the time-dependent receiver operating characteristic curve (AUC, *middle*), and Hazard Ratio (HR, *bottom*) estimated by the Kaplan Meier (KM) analysis of relapse-free survival (RFS) were used to compare the performances of the methods, respectively. The diamond in each box indicates the median value.

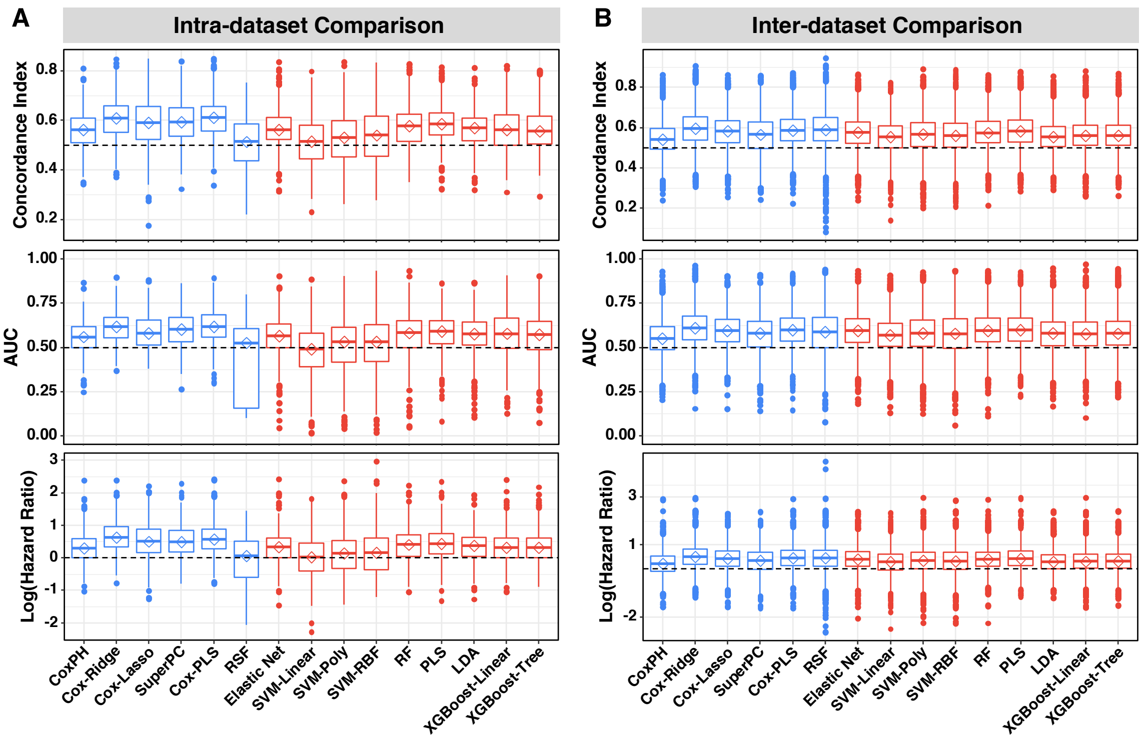


**Supplementary Figure S3. Comparison of survival analysis models versus binary classification models based on the same sample size for each dataset.** Prognostic performances of the machine learning (ML) algorithms were evaluated across all the 30 gene expression signatures across all the transcriptomics datasets with the censored data (no BCR occurred within 5 years following primary treatment) removed. Both intra-dataset (**A**) and inter-dataset (**B**) comparisons were performed. Median values of the three metrics, including Concordance index (C-index, *top*), area under the time-dependent receiver operating characteristic curve (AUC, *middle*), and Hazard Ratio (HR, *bottom*) estimated by the Kaplan Meier (KM) analysis of relapse-free survival (RFS) were used to compare the algorithms, respectively. The diamond in each box indicates the median value. Different colors of the box plots denote different categories of ML algorithms, *i.e.*, blue for the six survival analysis methods and red for the nine binary classification algorithms.

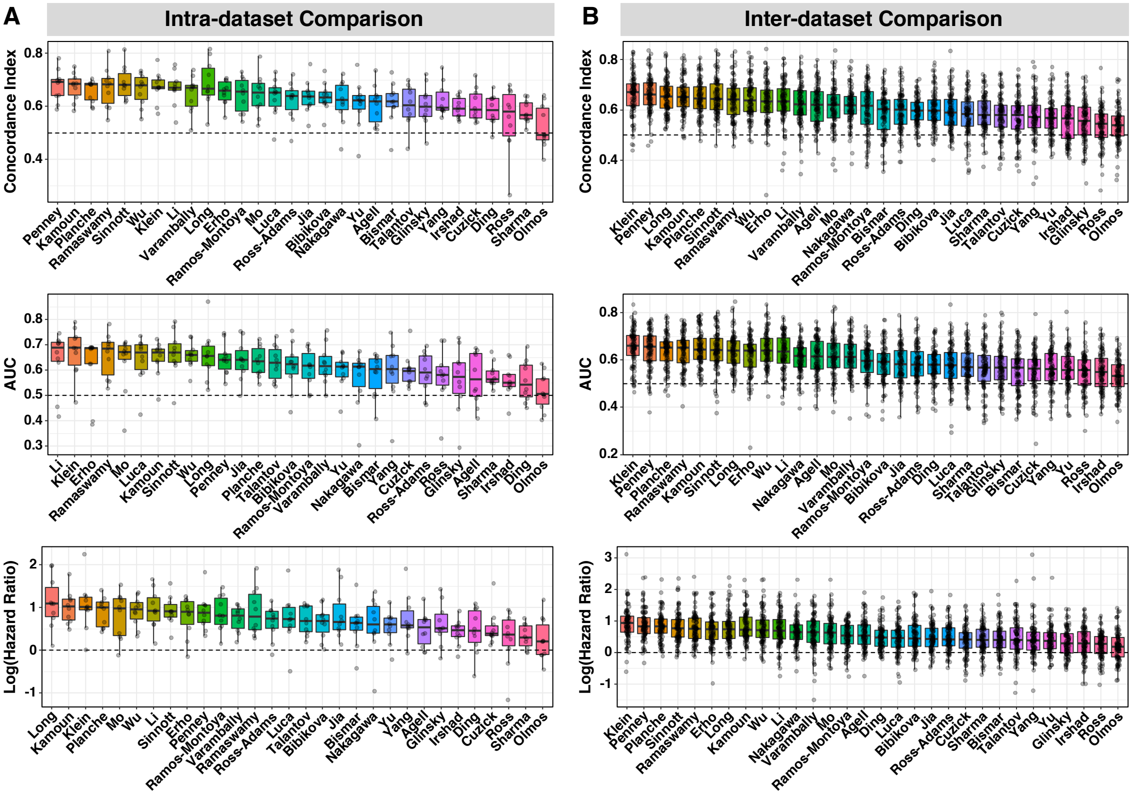


**Supplementary Figure S4. Evaluation of the 30 gene expression prognostic signatures based on Cox-PLS algorithm.** (**A**) Intra-dataset comparison via 10-fold cross-validation. (**B**) Inter-dataset comparison using one dataset as the training set and the remaining datasets as test sets. The prognostic signatures are ranked based on the median values of the three metrics, including Concordance index (C-index, *top*), area under the time-dependent receiver operating characteristic curve (AUC, *middle*), and Hazard Ratio (HR, *bottom*) estimated by the Kaplan Meier (KM) analysis of relapse-free survival (RFS), respectively, in the plots. Each dot represents the C-index, AUC, or HR value calculated within the dataset via 10-fold cross-validation for the intra-dataset comparison or in a test dataset for the inter-dataset comparison.

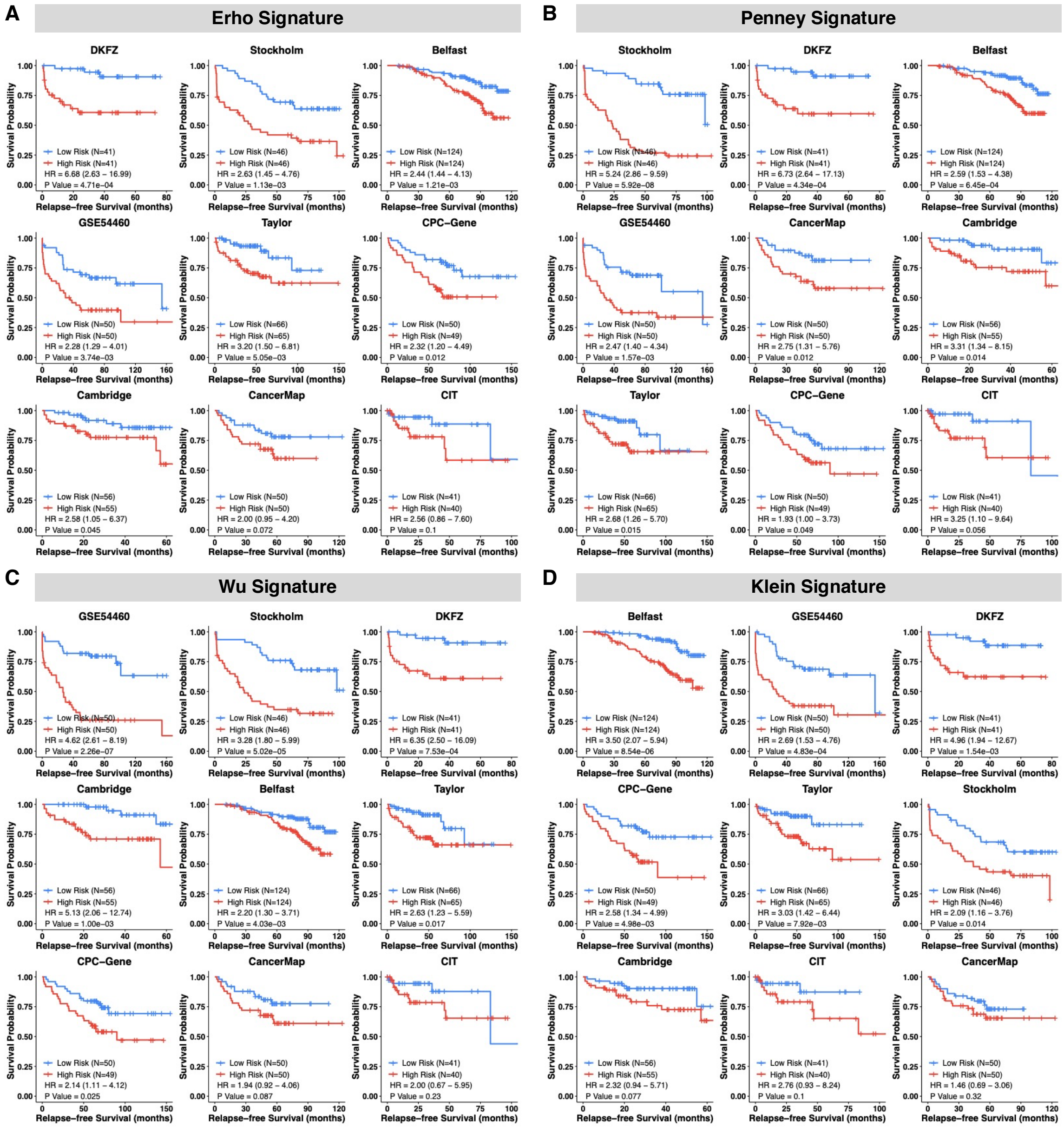


**Supplementary Figure S5. Kaplan Meier (KM) survival curves for some of the top prognostic signatures.** TCGA-PRAD cohort was used to train the prognostic models and the remaining nine independent cohorts were used to validate the models. Patients in each validation cohort were dichotomized into low- and high-risk groups based on the median value of the risk scores for KM analysis of relapse-free survival (RFS).

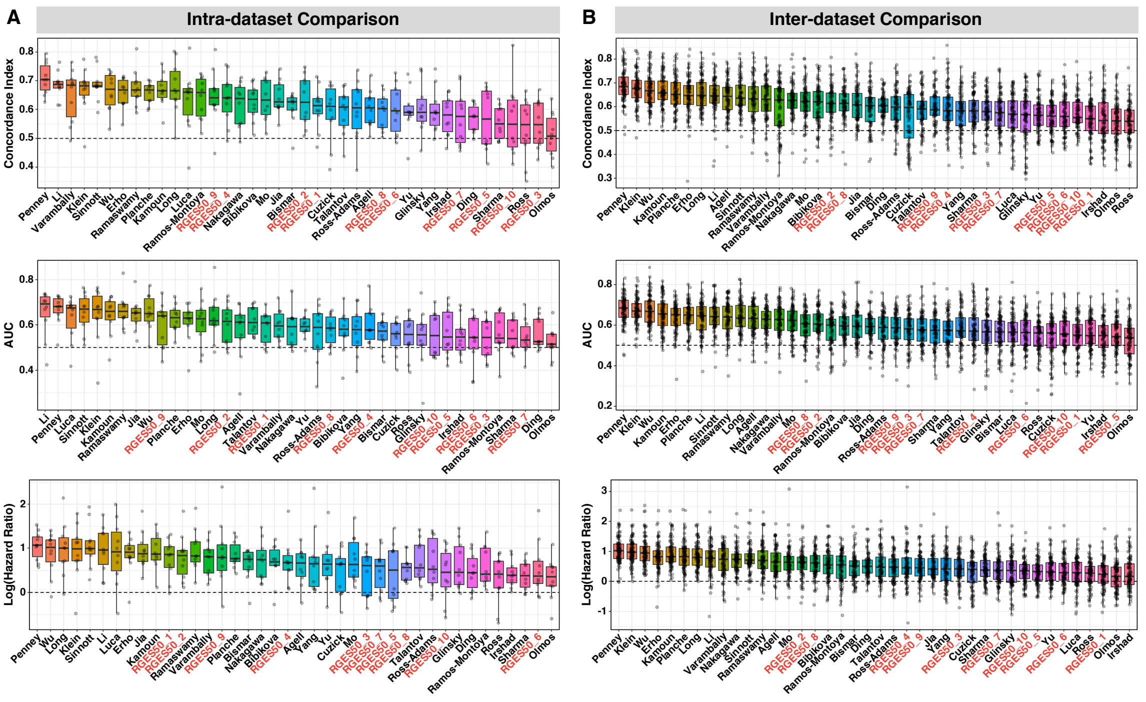


**Supplementary Figure S6. Comparison of the 30 published gene expression signatures with 10 random gene expression sets (RGESs) for prostate cancer prognosis based on the robust Cox-Ridge algorithm.** (**A**) Intra-dataset comparison via 10-fold cross-validation. (**B**) Inter-dataset comparison using one dataset as the training set and the remaining datasets as test sets. The published prognostic signatures and RGESs are ranked based on the median values of the three metrics, including Concordance index (C-index, *top*), area under the time-dependent receiver operating characteristic curve (AUC, *middle*), and Hazard Ratio (HR, *bottom*) estimated by the Kaplan Meier (KM) analysis of relapse-free survival (RFS), respectively, in the plots. Each dot represents the C-index, AUC, or HR value calculated within the dataset via 10-fold cross-validation for the intra-dataset comparison or in a test dataset for the inter-dataset comparison. The 10 RGESs are labeled in red.

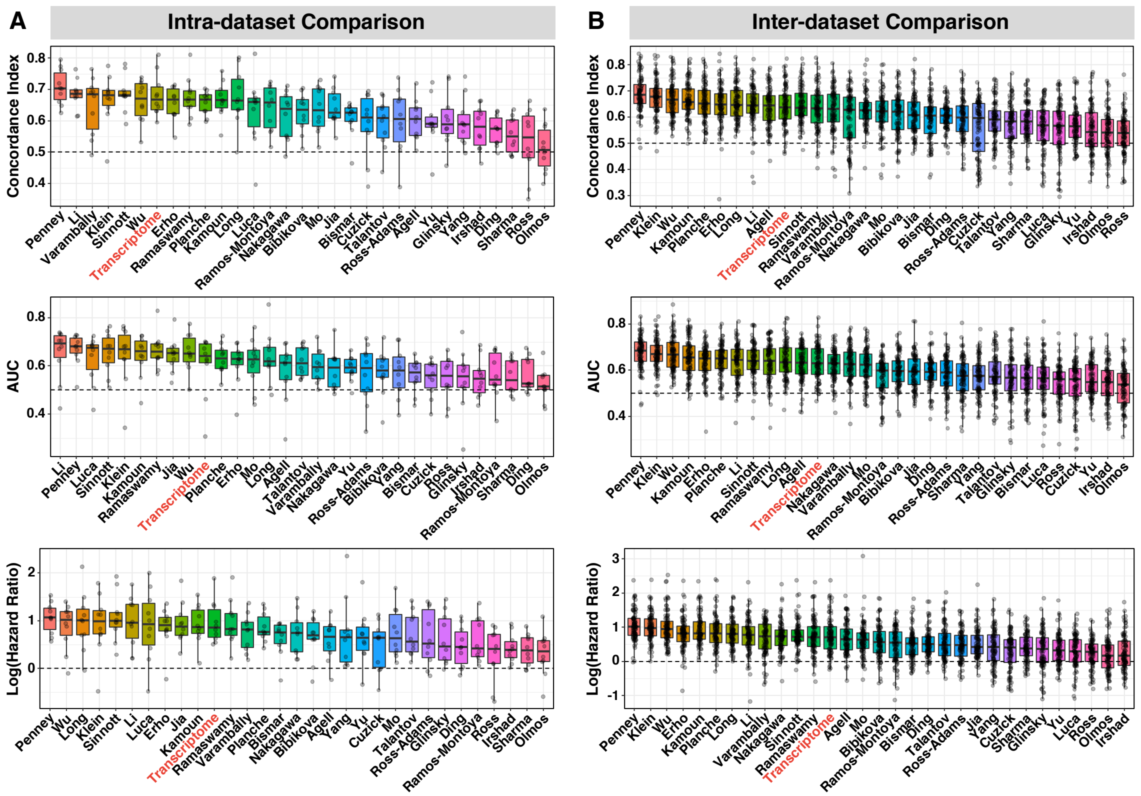


**Supplementary Figure S7. Comparison of the 30 published gene expression signatures with the whole transcriptome for prostate cancer prognosis based on the robust Cox-Ridge algorithm.** (**A**) Intra-dataset comparison via 10-fold cross-validation. (**B**) Inter-dataset comparison using one dataset as the training set and the remaining datasets as test sets. The published prognostic signatures and the whole transcriptome signature are ranked based on the median values of the three metrics, including Concordance index (C-index, *top*), area under the time-dependent receiver operating characteristic curve (AUC, *middle*), and Hazard Ratio (HR, *bottom*) estimated by the Kaplan Meier (KM) analysis of relapse-free survival (RFS), respectively, in the plots. Each dot represents the C-index, AUC, or HR value calculated within the dataset via 10-fold cross-validation for the intra-dataset comparison or in a test dataset for the inter-dataset comparison. The prognostic signature based on the whole transcriptome is labeled in red.

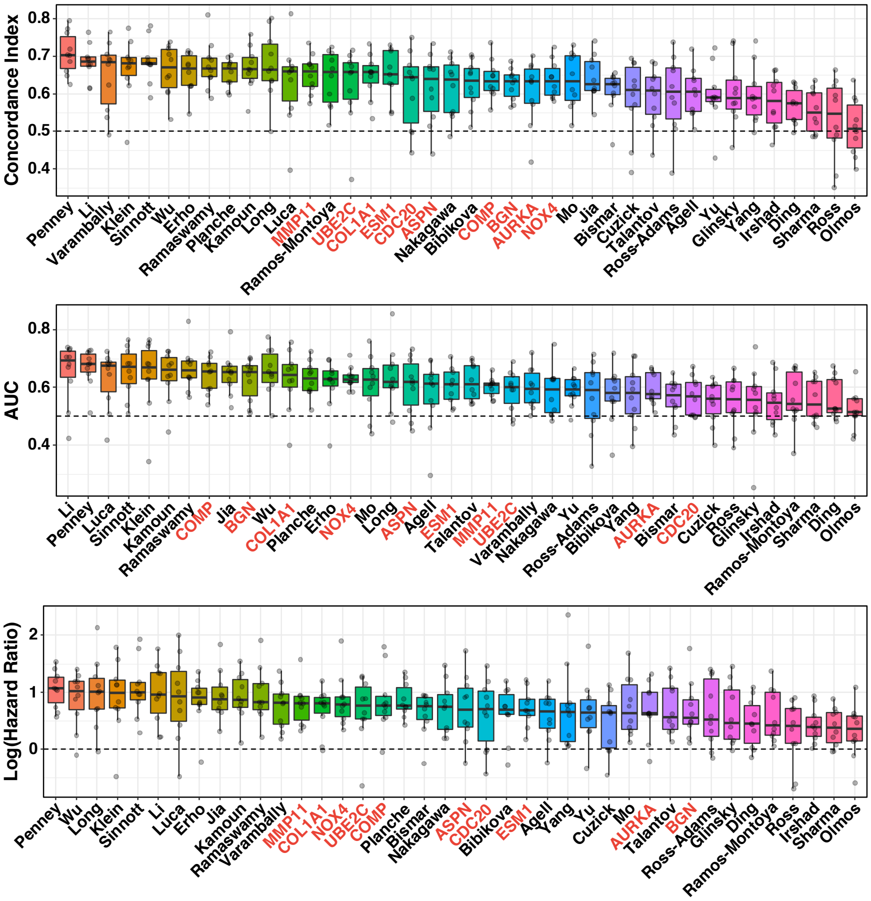


**Supplementary Figure S8. Comparison of the 30 published gene expression signatures with the top 10 individual genes for prostate cancer prognosis based on the robust Cox-Ridge algorithm.** For the 30 published signatures, 10-fold cross validation within each dataset was used to train the model and compute the risk score for each patient, whereas for the 1032 signature genes, the expression values were used directly, for the performance evaluation. Only the top 10 signature genes are included in each plot. The published prognostic signatures and the top 10 individual signature genes are ranked based on the median values of the three metrics, including Concordance index (C-index, *top*), area under the time-dependent receiver operating characteristic curve (AUC, *middle*), and Hazard Ratio (HR, *bottom*) estimated by the Kaplan Meier (KM) analysis of relapse-free survival (RFS), respectively, in the plots. Each dot represents the C-index, AUC, or HR value calculated within the dataset. The top 10 individual signature genes are labeled in red.

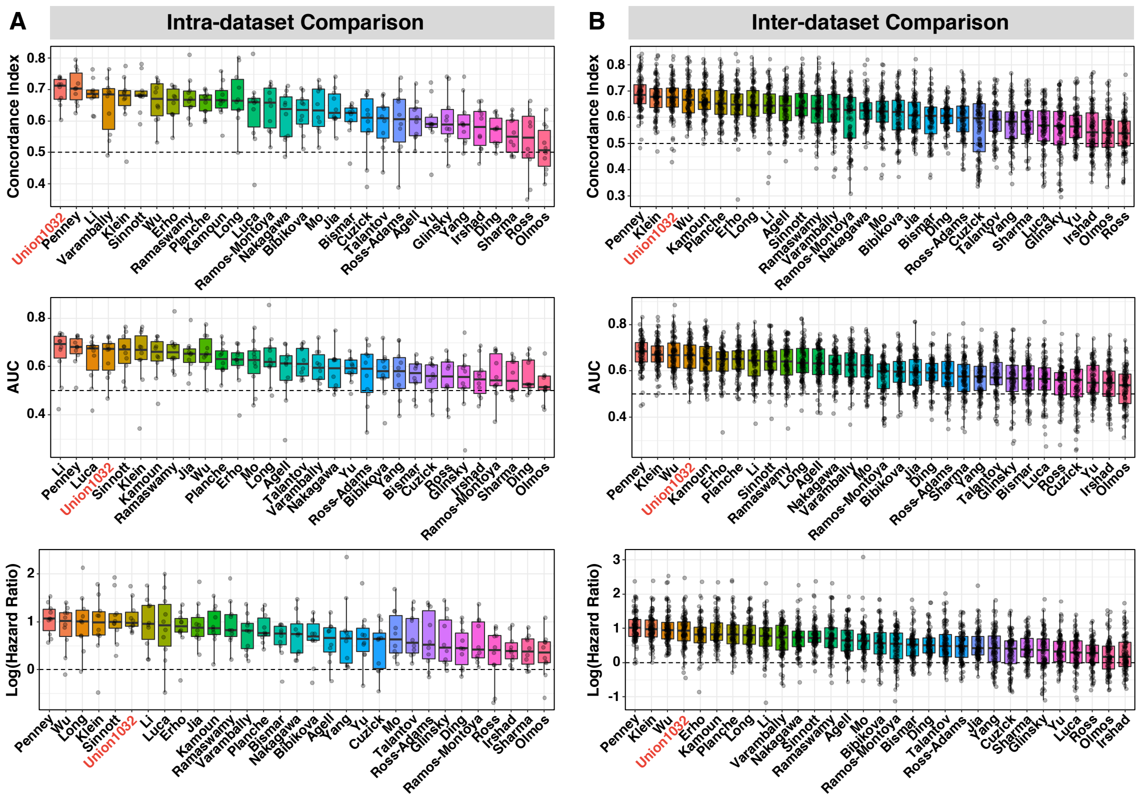


**Supplementary Figure S9. Comparison of the 30 published gene expression signatures with the union of 1032 signature genes (Union1032) for prostate cancer prognosis based on the robust Cox-Ridge algorithm.** (**A**) Intra-dataset comparison via 10-fold cross-validation. (**B**) Inter-dataset comparison using one dataset as the training set and the remaining datasets as test sets. The published prognostic signatures and the Union1032 signature are ranked based on the median values of the three metrics, including Concordance index (C-index, *top*), area under the time-dependent receiver operating characteristic curve (AUC, *middle*), and Hazard Ratio (HR, *bottom*) estimated by the Kaplan Meier (KM) analysis of relapse-free survival (RFS), respectively, in the plots. Each dot represents the C-index, AUC, or HR value calculated within the dataset via 10-fold cross-validation for the intra-dataset comparison or in a test dataset for the inter-dataset comparison. The Union1032 signature is labeled in red.

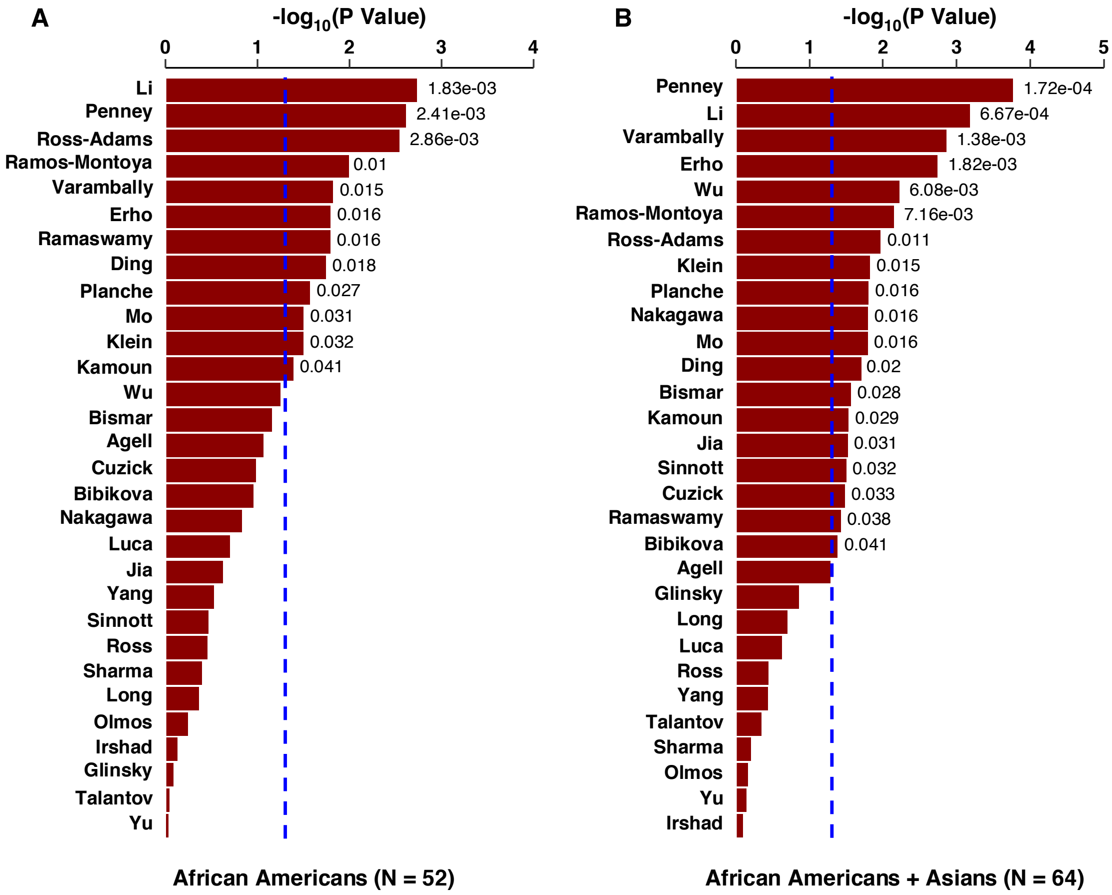


**Supplementary Figure S10. Validation of the 30 published signatures in non-white men.** The prognostic models were trained using the primary tumor samples from 386 white men in the TCGA-PRAD dataset based on the Cox-Ridge algorithm. Validations of the models were performed both in the cohort of 52 African Americans (A) and in the combined cohort of 64 African Americans and Asians (B) in TCGA-PRAD. Patients in each validation cohort were dichotomized into low- and high-risk groups based on the median value of the risk scores for Kaplan Meier (KM) analysis of relapse-free survival (RFS). Among the 30 published prognostic signatures, 12 of them can be validated in the African American cohort, and 19 signatures can be validated in the combined cohort of African Americans and Asians with p values less than 0.05.
