## Supplementary Table S2 for "Comprehensive evaluation of machine learning models and gene expression signatures for prostate cancer prognosis using large population cohorts"

**Supplementary Table S2.** Gene expression signatures for prostate cancer prognosis

| Signature Name | Alias | No. of Genes | Year | Citation |
| --- | --- | --- | --- | --- |
| Agell |  | 12 | 2012 | (1) |
| Bibikova |  | 16 | 2007 | (2) |
| Bismar |  | 12 | 2006 | (3) |
| Cuzick | Prolaris | 31 | 2011 | (4) |
| Ding |  | 4 | 2011 | (5) |
| Erho | Decipher | 19 | 2013 | (6) |
| Glinsky |  | 11 | 2005 | (7) |
| Irshad |  | 19 | 2013 | (8) |
| Jia |  | 15 | 2016 | (9) |
| Kamoun | CIT36 | 36 | 2018 | (10) |
| Klein | OncotypeDX | 12 | 2014 | (11) |
| Li |  | 160 | 2020 | (12) |
| Long |  | 24 | 2014 | (13) |
| Luca | DESNT | 45 | 2018 | (14) |
| Mo | SDMS | 93 | 2018 | (15) |
| Nakagawa |  | 17 | 2008 | (16) |
| Olmos | LPD1 | 9 | 2012 | (17) |
| Penney |  | 157 | 2011 | (18) |
| Planche |  | 36 | 2011 | (19) |
| Ramaswamy |  | 17 | 2003 | (20) |
| Ramos-Montoya |  | 222 | 2014 | (21) |
| Ross |  | 6 | 2012 | (22) |
| Ross-Adams |  | 100 | 2015 | (23) |
| Sharma |  | 16 | 2013 | (24) |
| Sinnott |  | 30 | 2017 | (25) |
| Talantov |  | 3 | 2010 | (26) |
| Varambally |  | 44 | 2005 | (27) |
| Wu |  | 29 | 2013 | (28) |
| Yang |  | 28 | 2018 | (29) |
| Yu |  | 14 | 2007 | (30) |

**References**

1.	Agell L, Hernández S, Nonell L, Lorenzo M, Puigdecanet E, de Muga S, et al. A 12-gene expression signature is associated with aggressive histological in prostate cancer: SEC14L1 and TCEB1 genes are potential markers of progression. Am J Pathol. 2012;181:1585–94.

2.	Bibikova M, Chudin E, Arsanjani A, Zhou L, Garcia EW, Modder J, et al. Expression signatures that correlated with Gleason score and relapse in prostate cancer. Genomics. 2007;89:666–72.

3.	Bismar TA, Demichelis F, Riva A, Kim R, Varambally S, He L, et al. Defining Aggressive Prostate Cancer Using a 12-Gene Model. Neoplasia. 2006;8:59–68.

4.	Cuzick J, Swanson GP, Fisher G, Brothman AR, Berney DM, Reid JE, et al. Prognostic value of an RNA expression signature derived from cell cycle proliferation genes in patients with prostate cancer: a retrospective study. Lancet Oncol. 2011;12:245–55.

5.	Ding Z, Wu C-J, Chu GC, Xiao Y, Ho D, Zhang J, et al. SMAD4-dependent barrier constrains prostate cancer growth and metastatic progression. Nature. 2011;470:269–73.

6.	Erho N, Crisan A, Vergara IA, Mitra AP, Ghadessi M, Buerki C, et al. Discovery and validation of a prostate cancer genomic classifier that predicts early metastasis following radical prostatectomy. PLoS One. 2013;8:e66855.

7.	Glinsky GV, Berezovska O, Glinskii AB. Microarray analysis identifies a death-from-cancer signature predicting therapy failure in patients with multiple types of cancer. J Clin Invest. American Society for Clinical Investigation; 2005;115:1503–21.

8.	Irshad S, Bansal M, Castillo-Martin M, Zheng T, Aytes A, Wenske S, et al. A Molecular Signature Predictive of Indolent Prostate Cancer. Science Translational Medicine. American Association for the Advancement of Science; 2013;5:202ra122-202ra122.

9.	Jia Z, Rahmatpanah FB, Chen X, Lernhardt W, Wang Y, Xia X-Q, et al. Expression Changes in the Stroma of Prostate Cancer Predict Subsequent Relapse. PLOS ONE. Public Library of Science; 2012;7:e41371.

10.	Kamoun A, Cancel-Tassin G, Fromont G, Elarouci N, Armenoult L, Ayadi M, et al. Comprehensive molecular classification of localized prostate adenocarcinoma reveals a tumour subtype predictive of non-aggressive disease. Annals of Oncology. Elsevier; 2018;29:1814–21.

11.	Klein EA, Cooperberg MR, Magi-Galluzzi C, Simko JP, Falzarano SM, Maddala T, et al. A 17-gene assay to predict prostate cancer aggressiveness in the context of Gleason grade heterogeneity, tumor multifocality, and biopsy undersampling. Eur Urol. 2014;66:550–60.

12.	Li R, Wang S, Cui Y, Qu H, Chater JM, Zhang L, et al. Extended application of genomic selection to screen multiomics data for prognostic signatures of prostate cancer. Brief Bioinform. 2021;22:bbaa197.

13.	Long Q, Xu J, Osunkoya AO, Sannigrahi S, Johnson BA, Zhou W, et al. Global transcriptome analysis of formalin-fixed prostate cancer specimens identifies biomarkers of disease recurrence. Cancer Res. 2014;74:3228–37.

14.	Luca B-A, Brewer DS, Edwards DR, Edwards S, Whitaker HC, Merson S, et al. DESNT: A Poor Prognosis Category of Human Prostate Cancer. Eur Urol Focus. 2018;4:842–50.

15.	Mo F, Lin D, Takhar M, Ramnarine VR, Dong X, Bell RH, et al. Stromal Gene Expression is Predictive for Metastatic Primary Prostate Cancer. Eur Urol. 2018;73:524–32.

16.	Nakagawa T, Kollmeyer TM, Morlan BW, Anderson SK, Bergstralh EJ, Davis BJ, et al. A Tissue Biomarker Panel Predicting Systemic Progression after PSA Recurrence Post-Definitive Prostate Cancer Therapy. PLOS ONE. Public Library of Science; 2008;3:e2318.

17.	Olmos D, Brewer D, Clark J, Danila DC, Parker C, Attard G, et al. Prognostic value of blood mRNA expression signatures in castration-resistant prostate cancer: a prospective, two-stage study. Lancet Oncol. 2012;13:1114–24.

18.	Penney KL, Sinnott JA, Fall K, Pawitan Y, Hoshida Y, Kraft P, et al. mRNA Expression Signature of Gleason Grade Predicts Lethal Prostate Cancer. JCO. Wolters Kluwer; 2011;29:2391–6.

19.	Planche A, Bacac M, Provero P, Fusco C, Delorenzi M, Stehle J-C, et al. Identification of Prognostic Molecular Features in the Reactive Stroma of Human Breast and Prostate Cancer. PLOS ONE. Public Library of Science; 2011;6:e18640.

20.	Ramaswamy S, Ross KN, Lander ES, Golub TR. A molecular signature of metastasis in primary solid tumors. Nat Genet. 2003;33:49–54.

21.	Ramos-Montoya A, Lamb AD, Russell R, Carroll T, Jurmeister S, Galeano-Dalmau N, et al. HES6 drives a critical AR transcriptional programme to induce castration-resistant prostate cancer through activation of an E2F1-mediated cell cycle network. EMBO Molecular Medicine. John Wiley & Sons, Ltd; 2014;6:651–61.

22.	Ross RW, Galsky MD, Scher HI, Magidson J, Wassmann K, Lee G-SM, et al. A whole-blood RNA transcript-based prognostic model in men with castration-resistant prostate cancer: a prospective study. Lancet Oncol. 2012;13:1105–13.

23.	Ross-Adams H, Lamb AD, Dunning MJ, Halim S, Lindberg J, Massie CM, et al. Integration of copy number and transcriptomics provides risk stratification in prostate cancer: A discovery and validation cohort study. EBioMedicine. 2015;2:1133–44.

24.	Sharma NL, Massie CE, Ramos-Montoya A, Zecchini V, Scott HE, Lamb AD, et al. The androgen receptor induces a distinct transcriptional program in castration-resistant prostate cancer in man. Cancer Cell. 2013;23:35–47.

25.	Sinnott JA, Peisch SF, Tyekucheva S, Gerke T, Lis R, Rider JR, et al. Prognostic Utility of a New mRNA Expression Signature of Gleason Score. Clin Cancer Res. 2017;23:81–7.

26.	Talantov D, Jatkoe TA, Böhm M, Zhang Y, Ferguson AM, Stricker PD, et al. Gene Based Prediction of Clinically Localized Prostate Cancer Progression After Radical Prostatectomy. The Journal of Urology. 2010;184:1521–8.

27.	Varambally S, Yu J, Laxman B, Rhodes DR, Mehra R, Tomlins SA, et al. Integrative genomic and proteomic analysis of prostate cancer reveals signatures of metastatic progression. Cancer Cell. 2005;8:393–406.

28.	Wu C-L, Schroeder BE, Ma X-J, Cutie CJ, Wu S, Salunga R, et al. Development and validation of a 32-gene prognostic index for prostate cancer progression. Proc Natl Acad Sci U S A. 2013;110:6121–6.

29.	Yang L, Roberts D, Takhar M, Erho N, Bibby BAS, Thiruthaneeswaran N, et al. Development and Validation of a 28-gene Hypoxia-related Prognostic Signature for Localized Prostate Cancer. EBioMedicine. 2018;31:182–9.

30.	Yu J, Yu J, Rhodes DR, Tomlins SA, Cao X, Chen G, et al. A Polycomb Repression Signature in Metastatic Prostate Cancer Predicts Cancer Outcome. Cancer Res. 2007;67:10657–63.
